# Probing stress-regulated ordering of the plant cortical microtubule array via a computational approach

**DOI:** 10.1101/2022.02.17.480928

**Authors:** Jing Li, Daniel Szymanski, Taeyoon Kim

## Abstract

Functional properties of cells, tissues, and organs rely on predictable growth outputs. A shape change in plant cells is determined by properties of a tough cell wall that deforms anisotropically in response to high turgor pressure. This morphogenesis requires tight coordination and feedback controls among cytoskeleton-dependent wall patterning, its material properties, and stresses in the wall. Cortical microtubules bias the mechanical anisotropy of cell wall by defining the trajectories of cellulose synthase motility as they polymerize bundled microfibrils in their wake. Cortical microtubules could locally align and orient relative to cell geometry; however, the means by this orientation occurs is not known. Correlations between the microtubule orientation, cell geometry, and predicted tensile forces are strongly established, and here we simulate how different attributes of tensile force can orient and pattern the microtubule array in the cortex. We implemented a discrete model with three-state transient microtubule behaviors influenced by local mechanical stress in order to probe the mechanisms of stress-dependent patterning. We varied the sensitivity of four types of dynamic behaviors observed on the plus ends of microtubules – growth, shrinkage, catastrophe, and rescue – to local stress and then evaluated the extent and rate of microtubule alignments in a square computational domain. We optimized constitutive relationships between local stress and the plus-end dynamics and employed a biomechanically well-characterized cell wall to analyze how stress can influence the density and orientation of microtubule arrays. Our multiscale modeling approaches predict that spatial variability in stress magnitude and anisotropy mediate mechanical feedback between the wall and organization of the cortical microtubule array.

**Author Summary:** Plant cell growth involve multiple steps and processes. During growth, cell shape changes continuously while responding to external cues from the surroundings. Since growth is mainly driven by pressure, mechanical properties of cell wall are crucial in regulating multiple biological processes that underlie cell expansion and growth. Cell wall assembly is dynamically coupled to the remodeling of subcellular proteins. Experimental evidence has confirmed there exists potential mechanical feedback between wall assembly and protein-protein interactions. However, the actual mechanism remains unknown. In this study, we develop a computational model to study how mechanical stress could affect subcellular protein dynamics or interactions and lead to their reorganization, reminiscent of continuous changes in global pattern and cell morphology. Our results identify key parameters that can respond to external mechanical stimuli at the cellular scale. We also show that a biological stress pattern could induce protein filament organization and bundles that mimic real subcellular structure from experimental images. These computational results could benefit design of experiments for studying and discovering the potential protein candidates that underlie the mechanical feedback between multiple cellular components. In this way, a more systematic understanding about plant cell growth could be achieved, with an integrated theory that combine biology, chemistry, mechanics, and genetics.

## Introduction

Tissue morphogenesis requires coordinated shape changes of isolated and adherent cells that are often driven by mechanics. For example, plant cells grow in a highly anisotropic fashion since turgor pressure expands the cell wall with anisotropic stiffness (1–3). Cellulose-dependent wall anisotropy underlies the morphological diversity in the plant kingdom (2, 4), and the orientation of the cellulose microfibrils is determined in large part by the cortical microtubule cytoskeleton. Cortical microtubules are tightly coupled to the plasma membrane (5) and bias the directions of active cellulose synthase (CESA) complexes (2); the cell wall becomes stiffer in a direction in which the cellulose fibers are synthesized and oriented, so elongation is restricted in that dimension. A major challenge in the field of plant development is to learn how morphologically potent cortical arrays are organized relative to cell and tissue geometries.

Microtubules are highly dynamic polymers in the cell, with individual microtubules and the entire microtubule network turning over on the time scale of minutes (6–8). The underlying dynamic instability of microtubules mediates these unpredictable behaviors (6, 7, 9, 10). Microtubules are cytoskeletal polymers comprised of α and β tubulins with polarity defined by plus and minus ends. The nucleation of microtubules in the plant cortex can occur de novo or via branching from pre-existing microtubules (11–13). Although both ends of microtubules exhibit a frequent switch between growth, shrinkage, and pause states (three-state dynamics) (14), the plus end is far more dynamically unstable than the minus end. When microtubules in the plant cortex grow from the plus end with the dynamic instability, they often encounter other microtubules. Since they are tightly coupled to the cell wall, the microtubules collide to each other, which can lead to different consequences. When the plus end of a microtubules makes contact to other microtubule with an angle below ∼40°, its polymerization is reoriented in a direction parallel to the other microtubule, which is called zippering. By contrast, collision with a contact angle greater than ∼40° leads to either crossover or a transition from growth to shrinkage called catastrophe. The collision-induced catastrophe and zippering are known to mediate ordered cortical microtubule arrays emerging at cellular scales. It was shown that a balance between steep-angle catastrophe and shallow-angle zippering are necessary and sufficient for promoting self-organization and parallel configuration of microtubules (9, 15). Regardless of collision, microtubules undergoing rapid shrinkage can transition to a growth state, which is called rescue.

Cellular scale orientations of microtubules are often claimed to be strongly correlated with aspects of cell geometry such as size or local curvature, but in the context of tissues, this is often not the case. Epidermal cells with similar shapes adopt multiple types of microtubule configurations on the outer cell cortex over time (16), and different faces of the cell can simultaneously display distinct orientations (8, 17). The constricted regions of lobed pavement cells are emphasized as a location for microtubule alignment (18, 19). In reality, there is only subtle enrichment of transfacial microtubules at subsets of cell indentations (20), and tensile force in the wall is likely to be a patterning element (8). Polarized trichoblasts like leaf hairs and cotton fibers present a simpler case since they lack neighboring cells and biomechanically distinct cell faces. During their anisotropic growth phases, a collar of transverse microtubules is present along the cell flank distal to the apical microtubule depletion zone (MDZ) (21–24). Strict thresholds for microtubule alignment and the MDZ mediate the simultaneous tapering and polarized elongation without cell swelling (25). Overall, the mechanisms that define the cellular scale orientations of microtubules in plant cells are still unknown.

Computational modeling of microtubule array alignment has provided clues about how interactions between microtubules and destabilizing boundary conditions can orient the cortical microtubule array with respect to cellular features. Dixit and Cyr first introduced a Monte Carlo simulation for cortical microtubule arrays and highlighted the importance of zippering and collision-induced catastrophe for mediating network alignment (9). A subsequent two-dimensional (2D) model that included stochastic microtubule behaviors showed that biased alignment of microtubule in the transverse direction could occur when two edges act as catastrophe-inducing boundaries. Such a boundary effect was also observed in a three-dimensional (3D) model where the end walls (top and bottom boundaries) in a cylindrical domain act as either the catastrophe-inducing or reflective boundary (26). Both types of the boundary condition induced transverse alignment even if collision-induced catastrophe between microtubules was disabled. A recent 3D model by Chakrabortty et al. showed that a catastrophe-inducing boundary and cell-face-specific microtubule-destabilizing surfaces can potently generate oriented microtubule arrays (19). However, the prevalence of catastrophe-inducing boundaries and the mechanistic underpinnings of cell-face-specific control of microtubule behaviors are not well established. For example, a threshold of 2.5 µm for a radius of curvature for microtubule destabilization has been observed in some epidermal cell types (27). However lobing pavement cells and tapering trichoblasts have curvatures that greatly exceed the curvature threshold for microtubule destabilization (8, 25). In addition, in epidermal cells, microtubules are often highly aligned along the anticlinal walls perpendicular to the cell edge (8, 28, 29). This alignment is orthogonal to the orientation that would be expected with a strong catastrophe-inducing boundary at the interface of the outer periclinal and anticlinal walls.

The observed anticlinal wall alignment of microtubules is strongly correlated with the magnitude and direction of tensile forces in the wall predicted by finite element (FE) modeling (8). During recent decades, a correlation between mechanical stress, self-organization of cortical microtubule array, and morphogenesis has been analyzed (30, 31). Numerous studies have shown that microtubules can orient in parallel to predicted patterns of cell wall tensile forces, and remodeling of microtubules often occurs gradually and continuously in response to perturbations that are likely to alter tensile forces (18, 32–36).

The ability of microtubules to sense tensile forces has been shown in the functional attributes of the mitotic spindle (37) and the enhanced polymerization of microtubules in the in vitro reconstitution experiments (38–40). However, molecular sensors that enable cortical microtubules in plant cells to sense the magnitude and direction of mechanical stress are still unknown. The most plausible model for the sensors is what Williamson proposed more than 30 years ago (4). He suggested the existence of load bearing elements in the wall that integrate wall stress/strain with microtubule behaviors.

Based on known interactions between tensile forces and microtubule dynamics described above, we created a discrete model with transient behaviors of microtubules influenced by local mechanical stress. By exploring a wide parametric space and imposing physiologically relevant stress patterns, we defined a plausible constitutive relationship between local stress and microtubule dynamics. We adopted stress field data imported from validated FE models to a 2D microtubule simulation domain. We found that spatial patterns of stress directionality, magnitude, and anisotropy mimicking stress distribution on realistic cell walls lead to microtubule arrays commonly observed near the cell apex. Results in our study provide a more systematic understanding of how microtubules are self-organized by responding to physiologically relevant mechanical stimuli.

## Methods

### Microtubule dynamics

Each microtubule is represented by serially connected individual segments whose length is 100 nm. The nucleation of new microtubules takes place at the rate of 10 μm^-2^·min^-1^ at a random location without dependence on existing microtubules. Following approaches used in a previous study (41), it is assumed that the dynamic behavior of the plus end of microtubules can switch between growth, shrinkage, and pause states (three-state dynamics), whereas their minus end always undergoes slow shrinkage at a constant rate (Fig. 1A).

**Figure 1.**
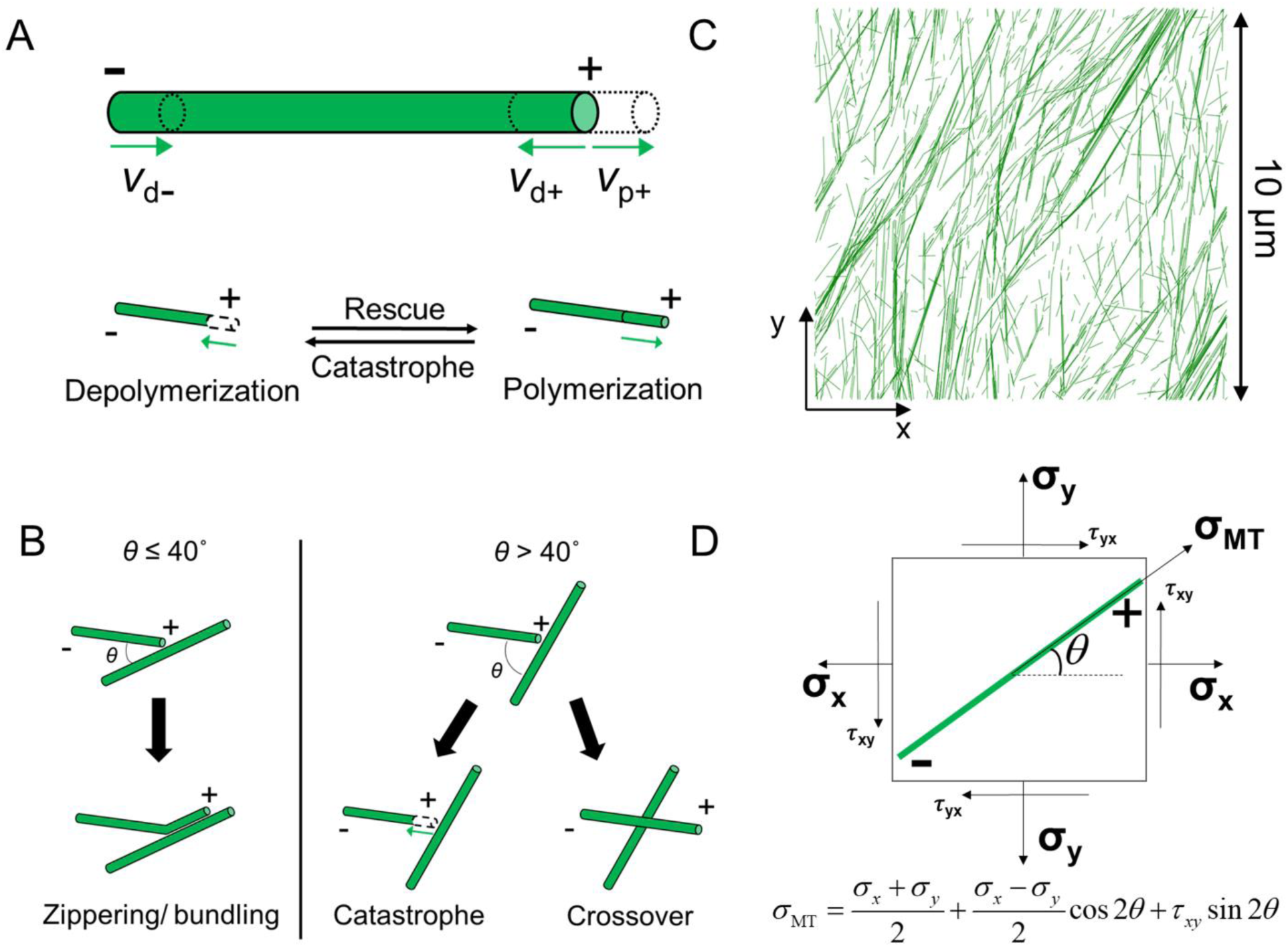
Model schematic and microtubule dynamics. (A and B) Schematic of MT dynamics, including stochastic events and angle-dependent collision/deterministic events. (A) Stochastic events include polymerization (v_p+_) and depolymerization (v_d+_) at the plus end and constant slow depolymerization (v_d-_) at the minus end, rescue, and catastrophe. (B) Angle-dependent collision-based events of MTs include zippering at a shallow angle (<40°) and catastrophe or crossover beyond a critical angle (>40°). (C) MT simulation domain is a 10 x 10 μm (initial domain size) square network with periodic boundary conditions in both x and y directions. MTs in the simulation domain are composed of serially connected small segments representing tubulin dimers, as shown in green. (D) Mathematical relationship between local principal stress and stress acting on individual microtubules depending on their angle of orientation.

Based on experimental observations, it is assumed that a collision between two microtubules with a contact angle smaller than 40° results in zippering meaning that one of the microtubules changes its orientation to align with the other microtubule (Fig. 1B). A spacing between microtubules is set to 25-50 nm which is close to observations in animal and plant cells (42, 43). A collision with a contact angle greater than 40° leads to either crossover or catastrophe. The probability of catastrophe after a collision is set to a value between 0.2 and 0.8 (41). Stochastic dynamic events of microtubules were updated once per 0.001 min, and collisions between microtubules were considered once per 0.05 min (Fig. S1A), which is consistent with the previous model (41).

The parameters used to depict the stochastic properties of microtubules and deterministic behaviors of microtubule-microtubule interactions are listed in Tables 1 and S1.

**Table 1.** List of microtubule dynamic parameters in the simulation for three-state models.

| Units<br>[min<sup>-1</sup>]<br>or<br>[μm/min] | WT(14) | Description |
| --- | --- | --- |
| $f_{gp}$ | 0.47 | Freq. growth to pause |
| $f_{gs}$ | 0.97 | Freq. growth to shorten |
| $f_{pg}$ | 0.51 | Freq. pause to growth |
| $f_{ps}$ | 0.24 | Freq. pause to shorten |
| $f_{sg}$ | 0.87 | Freq. shorten to growth |
| $f_{sp}$ | 0.23 | Freq. shorten to pause |
| $v_g^p$ | 3.69 | Growth rate (+ end) |
| $v_s^p$ | 5.80 | Shorten rate (+ end) |
| $v_s^m$ | 0.53* | Shorten rate (- end) |

\*: $0.253 (2.78 \text{ μm/min}) + 0.084 (-1.96 \text{ μm/min}) + 0.663 (0)$ Weighted average from (14).
Minus end growth phase (25.3%), pause phase (66.3%) and shorten phase (8.4%) considered in the weighted average.

### Stress pattern and computational domain

To investigate effects of local cell wall stress on microtubules, various stress patterns are mapped onto two types of computational domains: square and rectangular domains. For the square domain, we used 10×10 μm for its size with the periodic boundary condition (PBC) in x and y directions (Fig. 1C). We confirmed that this domain size is large enough to avoid finite-size effects by comparing results with those obtained from a larger domain with 20×20 μm in size (Figs. S2A, B). Stress patterns mapped onto the square matrix undergo either no, sudden, or gradual change during simulations. For the stress pattern with no change, stress components remain constant during the entire duration of simulations (100-200 min). For the stress pattern with a sudden change, an initial stress pattern is maintained for ∼100 min, and then a predominant stress direction is rotated by 90° and maintained till the end. For the stress pattern with a gradual change, stress components are varied linearly; the stress pattern is initially predominant in y direction, and the y component of stress is reduced linearly over time, whereas the x component is linearly increased. The x and y components of the stress become identical at the middle of these linear changes. For example, if it takes 50 min for the linear changes, stress becomes isotropic at 25 min. At the end, the extent of the stress anisotropy becomes identical to that of the initial stress anisotropy. It takes 50-150 min to complete the linear changes in the stress pattern, which is comparable to the timescales of tens of minutes or a few hours required for the pattern of cortical microtubules to be reoriented (44, 45).

For the rectangular domain, we used 30×10 μm for its size in the x and y directions. It is assumed that the longer direction (*x*) is parallel to the cell long axis, and the shorter direction (*y*) represents the transverse direction. In the stress pattern mapped onto the domain, the extent of stress anisotropy is relatively constant along the cell long axis, whereas the magnitudes of stresses decrease from left (the cell apex) to right (the cell base) to impose a stress gradient. The PBC is applied in the y direction to avoid finite-size effects. In the x direction, unless specified, there is no PBC to avoid an abrupt change in the stress across the boundaries. Instead, three different types of boundary conditions are imposed; the catastrophe-inducing boundary enforces growing microtubules to switch to a shrinkage state. The reflective boundary causes microtubules growing toward the boundary to change the direction. The repulsive boundary pauses a growing microtubule as soon as the microtubule makes contact to the boundary. These types of boundary conditions have been used in previous modeling studies (26, 41).

### Constitutive relationship between local stress and microtubule dynamics

The constitutive relationship between local stress and microtubules dynamics was devised from recent experiments showing that a tensile force up to ∼10 pN increases the polymerization rate and the rescue frequency but decreases the depolymerization rate and the catastrophe frequency (39). The range of stress on the cell wall that we estimated from the previous FE model is 3-15 MPa (25). If microtubules feel this large stress directly, the magnitude of forces from the stress is far beyond level that microtubules can sustain without rupturing (46). Thus, it is expected that microtubules are partially or indirectly coupled to the cell wall to sense the magnitude and direction of wall stress although it still remains unknown how stress is coupled to microtubules (4, 47).

Because microtubules align following the magnitude and direction of a tensile force, it is assumed that the magnitude of stress that a microtubule with a certain orientation affects one of the dynamics behaviors of microtubules (Fig. 1D). Using the stress transformation equation in continuum mechanics, we calculate the orientation-dependent local stress, *σ*_MT_ :

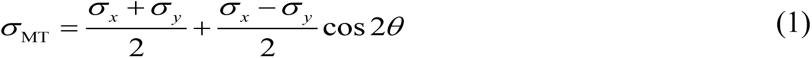

where *σ_x_* and *σ_y_* are normal stresses, and *θ* is the orientation of microtubules measured relative to the +x direction in the simulation space. Note that shear stress is assumed to be negligible in this equation, based on predictions from our previous FE model. A constitutive relationship was defined between *σ*_MT_ and either of the grow rate, the shrinkage rate, the catastrophe frequency, or the rescue frequency (Figs. S1B, C). We assumed either a linear, concave, or convex form for the constitutive relationship.

### Quantification of the order parameter and time constants

The order parameter (*S*_p_) is calculated to evaluate the extent of the ordering of microtubules:

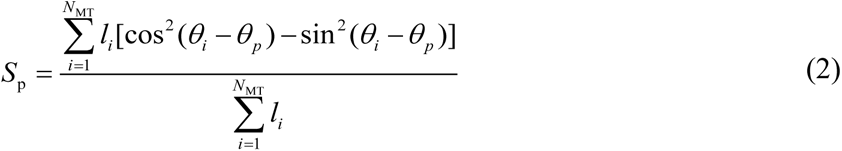

where *N*_MT_ is the number of microtubules, *θ_p_* is the dominant angle parallel to the direction of principal stress, and *θ_i_* and *l_i_* are the angle and length of each microtubule segment, respectively. We quantify a time constant to indicate the rate of a change in the order parameter. The first time constant (*τ*_p_) represents time at which *S*_p_ reaches half of its steady-state value. The second (*τ_p,_*_2_) and third (*τ_p,_*_3_) time constants indicate time at which *S*_p_ reaches 75% and 87.5% of the steady-state value, respectively. In addition, we also calculate the instantaneous rate of a change in *S*_p_.

### Live cell imaging

We performed live cell imaging for stage 4 trichome in which two to three short, blunt, branches were present. *Arabidopsis thaliana* seedlings were grown in Murashige and Skoog medium under continuous illumination. Single channel images were performed and collected with mCherry:MBD seedlings. Arabidopsis trichomes are imaged 10-12 DAG, where confocal fluorescence microscopy was performed using a Yokogawa spinning disk CSU-10 head mounted on a Zeiss Observer.Z1 inverted microscope. Stages 2-4 trichomes were imaged using a 100X PlanApo 1.46 oil-immersion objective while mCherry was excited by 561 nm laser lines. Minute-scale images were acquired by Evolve 52 camera (Photometrics) through band-pass filters (482.35 and 617/73; Semrock). The collected z-stacks were analyzed in ImageJ.

## Results

Our goal is to determine whether realistic stress patterns on the wall could act as a cellular-scale cue that can mediate microtubule array organization, using a computational model. In the model, microtubules are simulated as rigid polymers with polarity that exhibit dynamic behaviors determined based on live cell imaging data (14). As plant microtubules bind tightly to the plasma membrane (5), our 2D model can represent stochastic behaviors of microtubules and interactions between microtubules in the presence of the stress pattern. As shown in the flow chart (Fig. S1A), we implemented a stress pattern with various magnitudes and anisotropy with diverse assumptions for a constitutive relationship between local stress and microtubule dynamics (Fig. S1B). Then, we analyzed how the morphology of microtubule arrays changes depending on the assumption.

Tensile force patterns in cell wall are sensitive to the geometry of the cell or tissue organization (25, 36, 48, 49). Therefore, we tested if stress patterns obtained from the validated FE model of a polarized plant cell (25) could bias the cellular scale organization of the microtubule array. If the stress is isotropic, microtubule bundles align in random directions because microtubules are subjected to the same level of stress regardless of their orientations (Figs. 2A, B left and S3A, B left). This is the case, regardless of the size of domain (Figs. S2A, B). To evaluate the extent of microtubule alignment, the order parameter (*S*_p_) was calculated (Fig. 2C). *S*_p_ measured at a steady state is very similar to that in a case without any stress (∼0.4-0.5) and is not dependent highly on which dynamic parameter was affected by stress (Figs. 2D, S2C). As another measure to analyze the sensitivity of different microtubule parameters to stress, the half-time to reach steady state in *S*_p_ (*τ*_p_) was defined (Fig. 2C). *τ*_p_ did not vary significantly by different conditions when the four parameters were varied in response to an isotropic stress (Fig. 2E). The lifetime and length of microtubules were not enhanced by isotropic stress, compared to the case without stress (Table 2). When anisotropic stress with a 10:1 ratio (*σ*_x_ < *σ*_y_) was imposed, microtubules were aligned in the principal direction of stress to different extents depending on which parameter was modulated by stress (Figs. 2A, B right and S3A, B center). In all cases, *S*_p_ increased fast at early time points and slowed down later (Fig. S3C). Oriented stress increased *S*_p_ for all of the plus end variables compared to the isotropic stress control. However, the largest increase in *S*_p_ was observed when stress enhanced the polymerization rate or reduced the catastrophe frequency (Fig. 2D). Stress-dependent modulation of the depolymerization rate was the least potent in terms of effects on *S*_p_. *τ*_p_ did not differ significantly when the depolymerization rate and the rescue frequency were modulated by stress (Figs. 2E and S3E, F). However, *τ*_p_ was significantly reduced in both cases with the polymerization rate and the catastrophe frequency varied by stress. The average length and lifetime of microtubules were also enhanced more in those cases (Fig. S3D and Table 2). Larger changes *S*_p_, *τ*_p_, and the length and lifetime of microtubules predict that the polymerization rate and the catastrophe frequency were more efficient stress-sensitive regulators of microtubule alignment.

**Figure 2.**
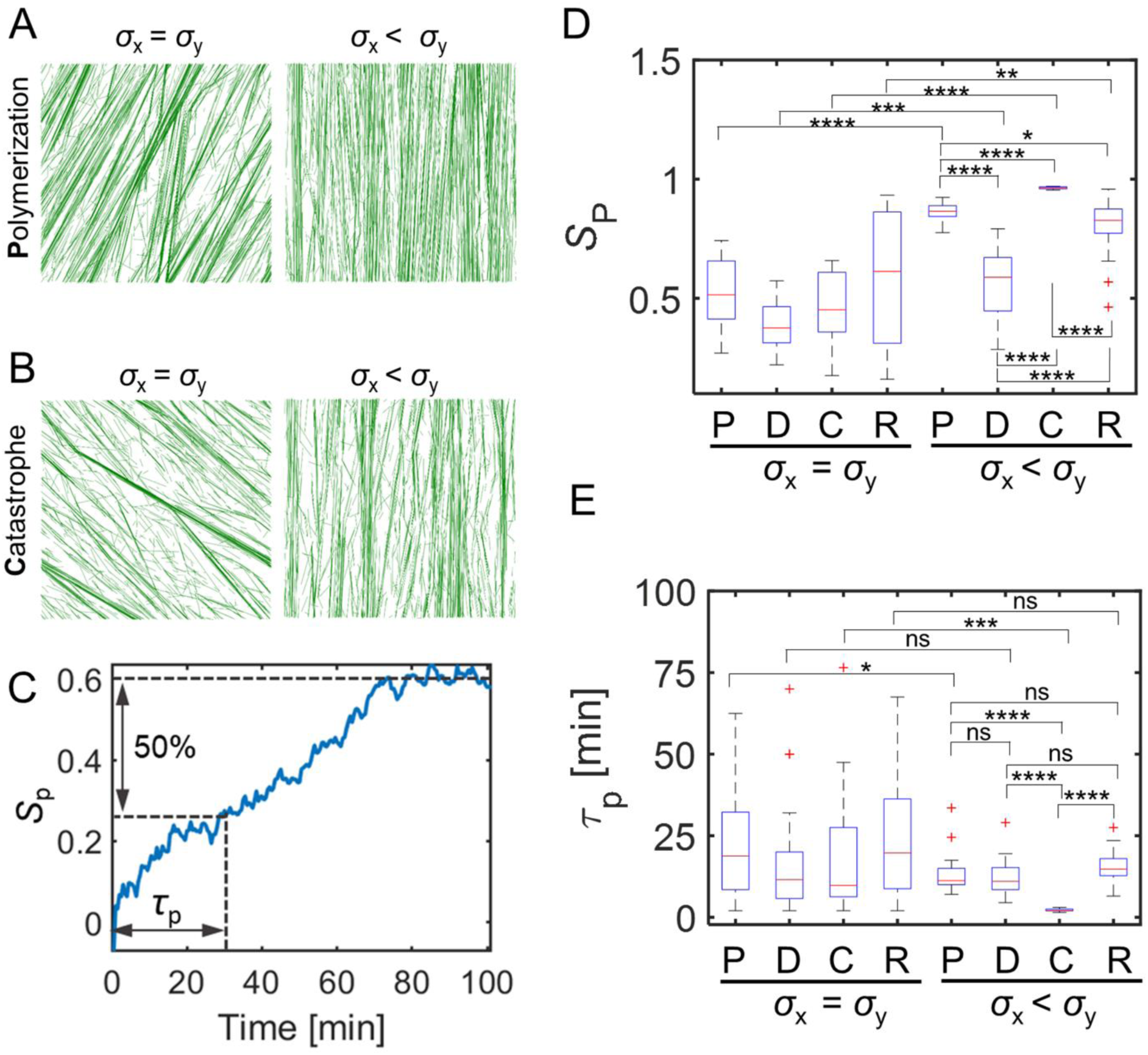
Anisotropic stress affects self-organization of microtubules and correlates with global orientation. (A and B) Steady state MT network morphology. All snapshots are taken at 100 min. MTs are subject to network stress predominant in y direction (right), isotropic (left). In A and B, polymerization rate is enhanced, and catastrophe frequency is reduced in alignment with principal stress, respectively. (C) A representative case with the time evolution of the network order parameter *S*_p_. The first time constant is calculated as the time required to reach half of the *S*_p_ value at steady state. (D) Summary of the network order parameter *S*_p_ at 100 min for all different conditions with isotropic vs. anisotropic stress in which principal stress influences individual stochastic parameter independently. (E) Boxplot of the time constants acquired from cases with isotropic vs. anisotropic stress in all conditions. Data for each condition are averaged over 20 simulations in each case (n=20). P: polymerization, D: depolymerization, C: catastrophe, R: rescue; ns p>0.05, * p<0.05, ** p<0.01, *** p<0.001, **** p<0.0001.

**Table 2.** Average microtubule lifetime and length under various stress conditions.

| $\tau_{\text{life}}$ [min]<br>$L_{\text{MT}}$ [$\mu\text{m}$] | | P<br>Polymerization | D<br>Depolymerization | C<br>Catastrophe | R<br>Rescue |
| --- | --- | --- | --- | --- | --- |
| Stress free<br>control | $\tau_{\text{life}}$ | 2.2 min | | | |
| | $L_{\text{MT}}$ | 2.0 $\mu\text{m}$ | | | |
| Isotropic stress<br>$\sigma_x = \sigma_y$ | $\tau_{\text{life}}$ | 2.3 min | 2.3 min | 2.3 min | 2.3 min |
| | $L_{\text{MT}}$ | 2.6 $\mu\text{m}$ | 2.0 $\mu\text{m}$ | 2.5 $\mu\text{m}$ | 2.2 $\mu\text{m}$ |
| Anisotropic<br>stress<br>$\sigma_x < \sigma_y$ or $\sigma_x > \sigma_y$ | $\tau_{\text{life}}$ | 2.9 min | 2.4 min | 3.0 min | 2.4 min |
| | $L_{\text{MT}}$ | 2.8 $\mu\text{m}$ | 2.2 $\mu\text{m}$ | 2.9 $\mu\text{m}$ | 2.3 $\mu\text{m}$ |

We next wanted to determine the threshold of stress anisotropy that is sufficiently high for increasing microtubule ordering. By testing different levels of stress anisotropy (*σ*_y_ / *σ*_x_), we found that the minimal ratio of 1.5:1 was required to induce microtubule alignment in the direction of principal stress (Fig. S4). In response to stress anisotropy with the minimal ratio, both *S*_p_ (Figs. S4A, C, E, G) and τ_p_ (Figs. S4B, D, F, H) exhibited a substantial increase or decrease, respectively (p < 0.001, n = 10). Previously, the validated FE model of a growing trichome branch showed that the estimated anisotropy of stress is ∼ 2:1 (25). This implies that the physiological level of the stress anisotropy is large enough to result in microtubule alignment, which is consistent with our simulation results.

It is not known how wall stress is coupled to factors that potentially modulate microtubule behaviors. It is possible that the responses of microtubules to stress can be non-linear, which could be attributed to many possible forms of cooperativity within a signal transduction cascade. Therefore, the constitutive relationship was varied from a linear function to a function with concave or convex shape to probe the sensitivity of the microtubule behaviors to a change in the relationship (Fig. S5). We either enhanced (+) or suppressed (-) the rate above a reference value. We did not observe a significant increase in the microtubule alignment when a concave function was used for the constitutive relationship (Fig. S1B, dash-dotted lines), compared to results obtained with the linear function (Figs. S5A, C). However, when a convex function was used (Fig. S1B, dashed lines), the efficiency of microtubule alignment is notably lower due to a relatively larger proportion of unstable microtubules although microtubules formed bundles in various orientations (Figs. S5B, D). This is consistent with the assumptions for microtubule dynamics (Fig. S1B).

In sum, our results demonstrated that microtubule alignment can emerge at the scale of the stress field if the directionality of a stress field affects stochastic dynamic behaviors occurring at the plus end of microtubules. However, we observed that there is a difference in the extent of microtubule alignment in the patterns, depending on which dynamic parameter is affected. It is also hard to neglect the effect by collision-induced catastrophe as it contributes to the turnover of microtubules. Therefore, we explored a 2D parametric space including stress anisotropy and collision-induced catastrophe. We found that stress anisotropy and collision-induced catastrophe can coregulate microtubule ordering (Supplementary Text and Fig. S6).

### Stress reorientation can reorganize aligned microtubule patterns

Tensile forces that influence microtubule organization are affected by tissue-level tension and the growth states of underlying cells (33, 50). Therefore, stress patterns involved with the tensile forces are not static features but can change at different time scales. As such, dynamic stress patterns can lead to the reorientation of cortical microtubule array (25, 34). To determine whether an established ordered microtubule array can be reoriented by a change in stress anisotropy, we imposed a dynamic stress pattern that is dominant in y direction at the beginning for 100 min and then undergoes a directional shift at the mid of the simulation. We assumed that the catastrophe frequency is affected by the time-varying stress because it was found to be the most efficient stress-sensitive parameter for microtubule alignment. A transition in the stress pattern occurred either instantaneously or gradually to reflect the slow shape change of plant cells that occurs on timescales of tens of minutes. The ratio of stress was 2:1 both at the beginning and at end of the transition.

When the principal direction of stress was rotated by 90° instantaneously at ∼100 min, well-aligned microtubules were quickly depolymerized due to a reduction in their average lifetime compared to the value before 100 min (Fig. S7A). In addition, newly nucleated microtubules were polymerized more persistently and then stabilized in the direction parallel to the new principal stress. This is due to the abrupt change in the differentiated catastrophe frequency by the rotation of stress pattern. It took ∼5 min for aligned microtubules to be transformed into randomly oriented microtubules. 50 min after the abrupt stress reorientation, most of microtubules were reoriented in the perpendicular direction (Fig. 3C), followed by stabilization and thickening of bundles due to a decrease in collision-induced catastrophe events.

**Figure 3.**
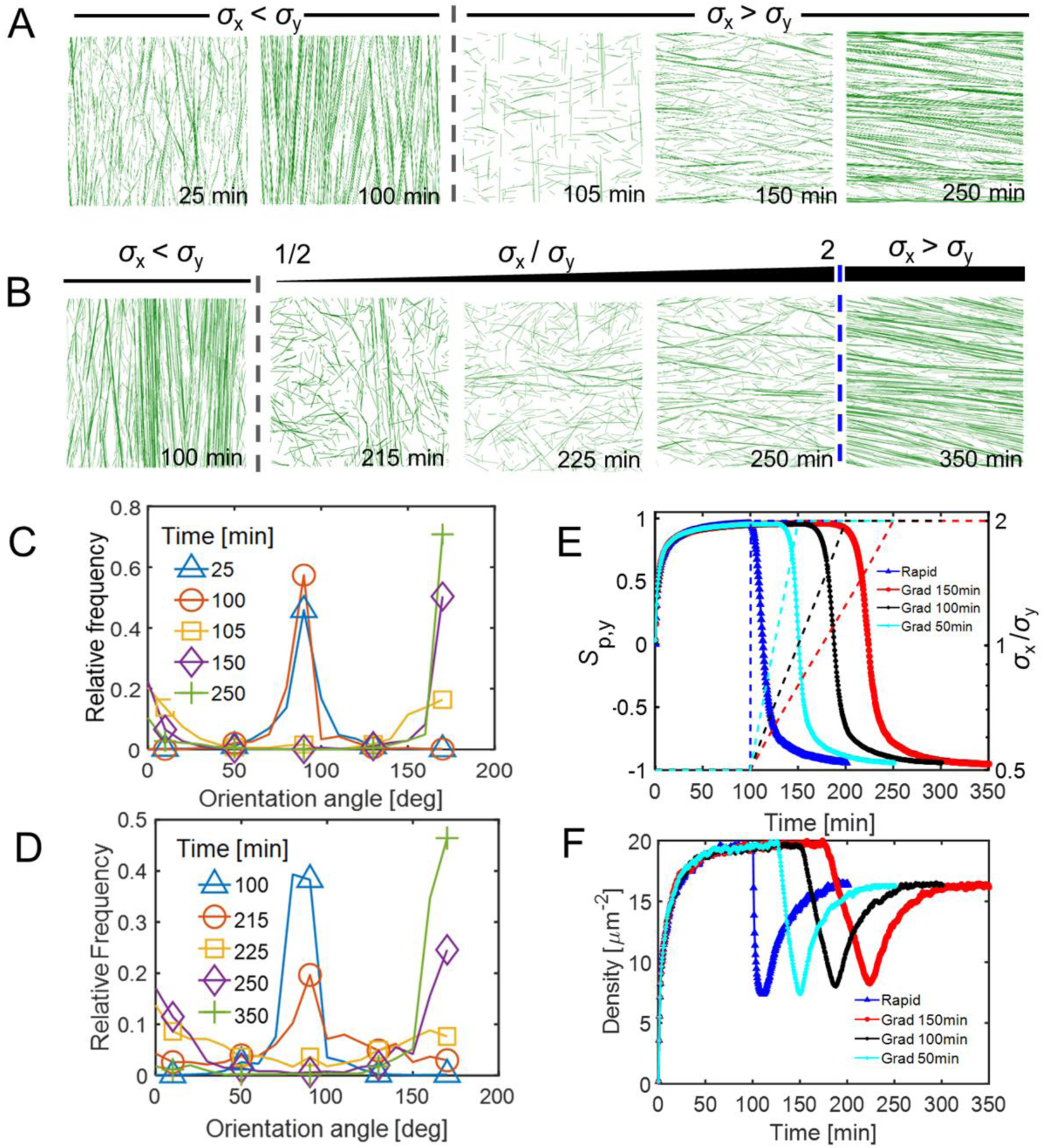
Dynamic stress pattern reorientation leads to remodeling of the microtubule network. (A and B) Time evolution of the MT network morphology (at various timepoints including 25, 100, 105, 150, 250 min for a rapid transition and 100, 215, 225, 250, 350 min for a gradual transition). Grey dashed line indicates the timepoint (100 min) after which reorientation of anisotropic stress takes place. The blue dash line indicates the timepoint after which anisotropic stress is stabilized. Initial network stress pattern is predominant in y direction. In A, stress pattern reorients instantaneously and becomes predominant in x direction after 100 min. In B, stress pattern gradually reorients until 250 min and becomes steady afterwards. (C and D) Relative frequency showing the distribution of microtubule orientation angle at various timepoints during stress pattern reorientation in A and B respectively. 90° is equivalent to the direction parallel to y axis, and 0° or 180° is equivalent to the direction parallel to x axis. A flat curve indicates homogeneous distribution of MT orientation angles. (E, F) Time evolution of the network order parameter *S*_p,y_ (E) and network microtubule segment densities (F) in four cases: rapid transition (as indicated in A, blue), gradual transition of stress (150 min, as indicated in B, red; 100 min, black; 50 min, cyan).

When the stress pattern was gradually changed, microtubules were initially aligned in the y direction perpendicular to the new stress as in the case with the instantaneous stress change (Fig. 3D). As stress anisotropy (*σ*_x_ / *σ*_y_) was enhanced over time (150 min), more short microtubule segments emerged in the x direction. At ∼215 min, the microtubule array became relatively homogeneous with randomly oriented microtubules (Figs. 3B, D). At 350 min, most of the microtubules were reoriented to the x direction, and bundles looked quite similar to those observed at the end of simulations run with the instantaneous stress change (Figs. 3A, B). The overall results were similar regardless of how long it took for the gradual stress change to take place. In the cases where it took 50 min or 150 min instead of 100 min, *S*_p_ showed an increase and reached similar level at the end (Fig. 3E). The complete reorientation, in all cases, took ∼50 min. In addition, it took ∼20 min for the network to become homogeneous (i.e., *S*_p_ ∼ 0) at a timepoint where stress became nearly isotropic (Fig. 3E). These values are similar to 10 min to 2 h which is a experimentally determined rate for microtubule reorientation (51–53). The average lifetime of microtubules aligned in various directions did not show a significant difference between cases with instantaneous and gradual stress changes (Figs. S7A, B). Local heterogeneity of microtubule orientations was positively correlated with the stress pattern transition (Fig. S7C). Interestingly, during microtubule orientation, we noticed microtubule density was almost halved when the microtubule array became homogeneous, followed by a rescue when the new bundles formed (Fig. 3F).

### Stress gradients generate fine-scale patterning of microtubule arrays

Tensile forces cannot be measured directly in plant cells and, in most cell types, stress patterns on cell walls are largely unknown. However, realistic wall stress patterns have been predicted for elongating leaf trichoblasts (Fig. 4A) based on known cell wall thickness gradient and a validated FE model (25). Cell wall stress was not constant along the cell surface. In young branches of this cell type, the microtubule array is highly organized, forming an apical MDZ and the extended collar of transverse microtubules that become progressively less dense toward the branch base (Fig. 4B) (23, 25). We tested whether spatially varying stress could be sufficient for generating this cell-scale patterns of microtubules.

**Figure 4.**
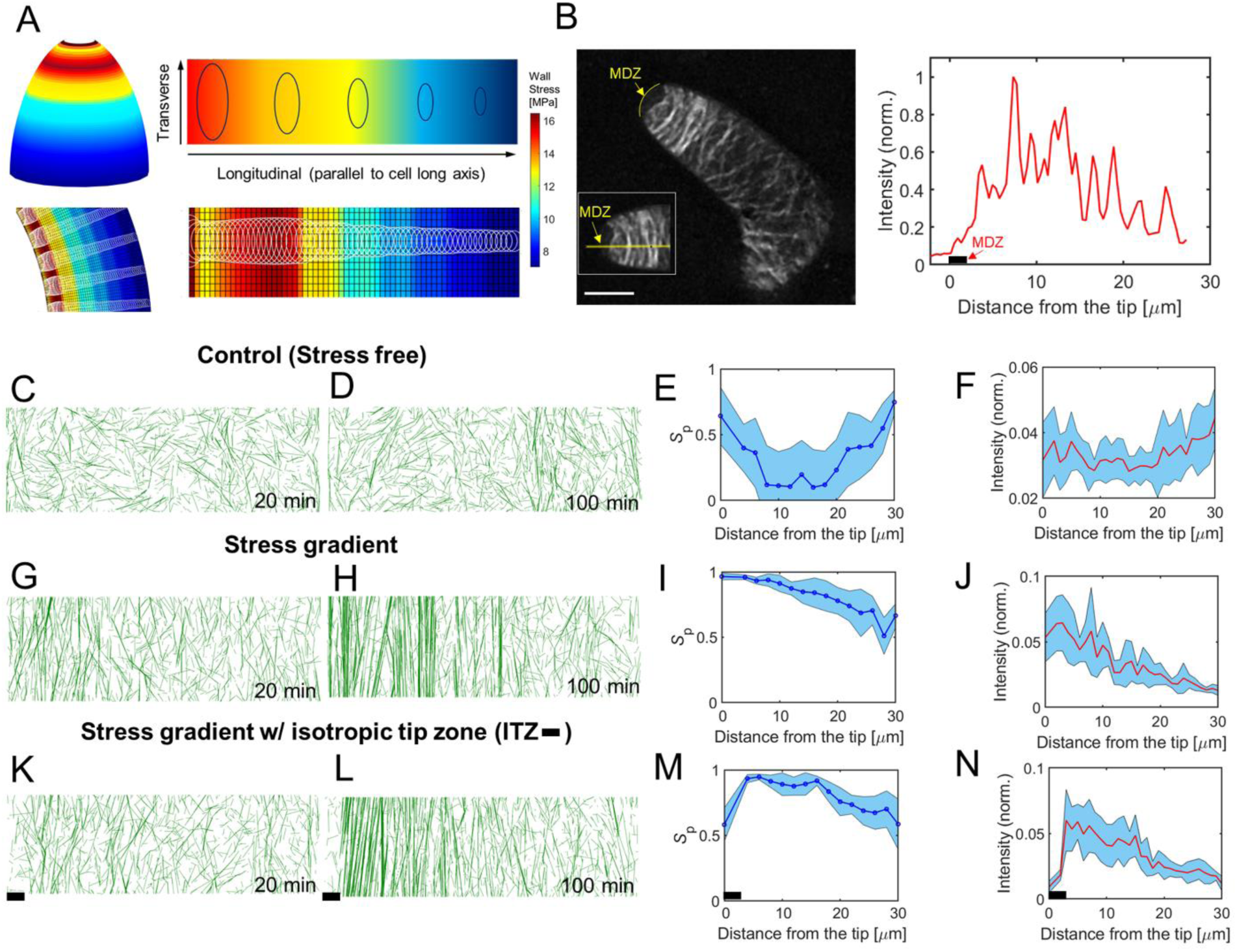
Physiological relevant stress gradient can induce distinguishable transverse microtubule bands. (A) Schematic of the stress pattern gradient pertinent to a finite element model of trichome morphogenesis. Magnitude of stress decreases from left to right, along the direction parallel to cell long axis while the stress is highly anisotropic and predominant in the transverse direction, indicated by the ellipses. The lower panels show an unfolded stress pattern based on the model results. The unfolded meshed stress pattern is mapped onto the simulation domain. (B) Representative confocal image showing microtubule localization in stage 4 young trichome branches (left panel). Transverse band is prominent near the cell apex. The microtubule depleted zone (MDZ) is highlighted and marked in yellow. Scale bar: 10 μm. Right panel shows the intensity profile from the apex to the base in the image shown in the left panel. (C-N) A catastrophe-inducing boundary condition is implemented on the two short edges of the simulation domain (30 μm x10 μm). Catastrophe-inducing boundary: when a growing microtubule encounters a boundary, it switches to a shrinkage state. The long edges are assumed to have periodic boundary conditions. MT band formation along the transverse direction is noticeable in cases with a stress gradient. (C, G, K) Early-time (20 min) and (D, H, L) steady-state (100 min) morphology of network in stress free (C, D), stress gradient (G, H), and stress gradient with isotropic stress in the tip zone (K, L, 2μm tip zone marked by black bar). (E, I, M) Local order parameter of microtubules in subregions of cases in D, H, L respectively. For this, the network was divided into fifteen subregions, incrementally separated by 2μm along the x direction. Blue curve indicates and shaded region represents standard deviation (n=10). (F, J, N) Normalized intensity of MT segments (tubulin dimers) as a function of the distance from the tip (left boundary), which is nearest to apex of a real cell. (F) For a homogenous network, the distribution of MT segments is relatively even along the cell long axis. (J) For a network with stress gradient, the distribution of MT segments peaks near the boundary and gradually decreases along the cell long axis. (N) With an isotropic stress zone near the tip (black bar), the distribution is similar to (J) except there is a decline in the intensity of microtubules in the zone. The distribution is consistent with the right panel in (B).

We adopted the stress pattern of the curved FE model of the branch for a rectangular simulation space of 30×10 μm in size (Fig. 4A). The stress pattern shows principal stress decreasing from the apical flanks of the tapered to the base, which is attributed to a combination of tapered morphology and an increasingly thick cell wall at the base (25). To analyze boundary effects, we imposed four different types of conditions on two boundaries normal to the longitudinal cell orientation: catastrophe-inducing, reflective, repulsive, and a periodic boundary as a control (Fig. S8A).

We first tested control cases with global isotropic stress or without stress in the presence of the periodic boundary or catastrophe-inducing boundary condition. Under stress-free condition with the periodic boundary condition in all directions, the microtubule array was homogeneous with small bundles, leading to local order parameter, *S*_p_(*z*), and local density, *I*(*z*), of microtubules that do not significantly vary in the z direction parallel to the longitudinal axis of the cell (Figs. S9A, B). Note that z = 0 corresponds to the location of the left boundary. When two catastrophe-inducing boundaries were employed, more aligned bundles aggregated near regions close to the two boundaries, characterized by an increase in *S*_p_(*z*) and *I*(*z*) (Figs. 4C-F and S9A, B). Under the isotropic stress condition, *S*_p_(*z*) showed a qualitatively similar distribution. However, isotropic stress significantly enhanced *S*_p_(*z*) globally by ∼0.3 due to increased average length and lifetime of microtubules (Table 2). These results indicate that catastrophe-inducing boundary alone could only increase local heterogeneity without a substantial change in global alignment. Next, we implemented a stress gradient. We observed clustering of microtubules at the tip (i.e., left boundary), regardless of the type of imposed boundary conditions (Fig. S8F). With catastrophe-inducing, reflective and repulsive boundaries, bundles were formed in the direction parallel to principal stress (i.e., perpendicular to the cell axis) in regions with large stress (Figs. 4G-J and S8B-E). *S*_p_(*z*) did not show a clear difference compared to the control case (Figs. 4E, I). Regions with higher stress showed *S*_p_(*z*) greater than 0.6 within 20 μm from the distal boundary. By contrast, in regions proximal to the base region with lower stress, microtubules were noticeably less dense and more homogenous in all cases. This is attributed to a fraction of disordered microtubules that turn over quickly due to collision-induced catastrophe. To test whether in vivo stress patterns at the cell apex could generate MDZ, we incorporated a small apical zone of 2 μm in length with isotropic stress (*σ*_x_ / *σ*_y_ =1) at the left boundary. Note that the isotropic stress is also present in the FE model of apex (Figs. S10), and isotropic stress is a general feature of hemispherical domes. Interestingly, the apical isotropic stress zone resulted in a very localized sparse, disordered population of short microtubules similar to the base (Figs. 4K-N), with a sharp decrease in *S*_p_(*z*) near the tip zone. The rest of the domain showed a similar pattern to that observed with an intact stress gradient. The relative intensity of microtubules showed consistent results with the microtubule morphology. Thus, the combination of spatially varying gradients of stress magnitude and anisotropy could generate a microtubule pattern that closely resembles the cortical array observed in living cells (Fig. 4B).

## Discussion

The microtubule-microfibril coalignment module generates an anisotropic cell wall that has enabled the morphological diversification in the plant kingdom (1, 54). Although the microtubule array is highly dynamic and spatially variable, some cell types (21, 23, 55) and cell faces, such as the anticlinal walls of epidermal cells (8, 29), display highly aligned microtubules with the cell axis. The aligned microtubules can pattern parallel microfibrils and dictate polarized cell expansion. Computational simulations have provided a critical insight into how deterministic events associated with collisions between microtubules can influence array organization (9, 41, 56). The critical question of how microtubule arrays acquire alignment with cell geometry remains poorly understood. It was shown that catastrophe-inducing boundaries can lead to the organization of the cortical array (26, 27, 41). However, the extent to which geometrically-induced or biochemically-induced catastrophe operates in plants is not known. Tensile forces, which reflect geometric and compositional parameters relevant to persistent growth patterns, are likely to potently affect the alignment of cortical microtubules (8, 18, 34, 53).

In this study, we developed a computational framework to analyze the organization of microtubule arrays with an assumption that microtubules can sense the magnitude and direction of tensile force in the cell wall; it was assumed that direction-dependent stress affects one of stochastic behaviors that the plus end of microtubules exhibits. The temporal component of the model is based on experimentally determined stochastic properties (Table 1). Our simulations recapitulated physiologically relevant alignment and reorientation of microtubules that occur in cells and indicate that tensile forces can align the cortical microtubule array as a function of cell geometry. Specifically, we showed that anisotropic stress can align microtubules (Figs. 2A, B, D) and decrease the time required for the array to reach a steady state (Fig. 2E). We identified the most efficient stress-sensitive parameter that can lead to better microtubule alignment. Stress-dependent variation of the polymerization rate and catastrophe frequency was the most potent means to rapidly establish a aligned microtubule array aligned with anisotropic stress. Note that our assumption of an increased polymerization rate or a decreased catastrophe frequency due to tensile forces is consistent with results from in vitro studies that probed tensile-force effects on microtubule plus-end dynamics (39, 40). The plus ends of microtubules switch between growth, pause, and shrinkage states with dynamic instability, so a large fraction of short microtubules can disappear very easily. If stress increases the polymerization rate or reduces the catastrophe frequency of microtubules oriented in a certain direction, the aligned subpopulation of short new microtubules can grow fast by making polymerization more dominant. Suppression of the catastrophe events can break symmetry between polymerization and depolymerization better so that microtubules are aligned faster and to a greater extent.

In growing cells and tissues, the wall stress fields are constantly remodeled (albeit slowly) as the geometry and wall material properties change. Tensile forces and microtubule alignment can also change as a function of the growth dynamics of underlying tissues (33). We conducted the microtubule array simulations with time-varying stress patterns to measure the robustness of the system (Fig. 3). We found that the more stable states in the microtubule simulations were not predestined and fixed outcomes defined by the input variables. We observed that during microtubule reorientation, bundles with different orientations could coexist with rather homogeneous morphology. This was previously observed in experiments where mechanical perturbation was employed to alter microtubule patterns (18, 35). Our simulation approach captured the behaviors of an unstable system that uses dynamic instability variables of microtubules coupled to tensile force in order to rearrange the array.

Cell wall stresses serve as useful multiscale patterning elements to modulate microtubule arrays because the stresses are non-uniform on a cell surface and depend on both geometric and cell wall compositions. For example, tensile forces are locally concentrated and aligned at the interface where unpaired outer periclinal walls pull upward on anticlinal walls, and microtubules at this location are highly coaligned (8, 29). The magnitude of cell wall stress is also inversely proportional to cell autonomous parameters like wall thickness. In isolated leaf trichoblasts, subcellular cell wall thickness gradients exist (25). We showed here that the spatial information of the direction and magnitude of stress can be decoded to generate an ordered microtubule array with a tip to base density gradient that mirrors that of living cells (Figs. 4B, G-J).

Living trichoblasts have an apical MDZ that may reflect an additional gradient in stress anisotropy. In domed, cylindrically-shaped trichoblasts, root hairs, pollen tubes, and moss protonema, stress will be more isotropic at the apex compared to the cell flanks. Based on geometry alone, the flanks of cylindrically-shaped cells are expected to have anisotropy ratios of ∼2:1. This value exceeds the threshold of 1.5:1 that was needed to order the microtubule network in simulations (Fig. S4). The apex dome is an important site for signal transduction and cytoskeletal patterning (51), and it has been hypothesized that ROP-dependent recruitment of catastrophe-inducing Kinesin13-related proteins was involved in maintaining the MDZ (57). However, our computational simulations showed that a combination of spatially defined gradients of stress anisotropy and magnitude can pattern a microtubule array with the basic alignment and density features that reflect the MDZ and the apical collar of cortical microtubules (Figs. 4B, K-N).

We probed the effects of periodic or catastrophe-inducing boundaries on microtubule alignment with isotropic stress or without stress. We observed that microtubule density and alignment patterns were not similar to the experimental observations if anisotropic stress is not present (Fig. S9 and Supplementary Text). These results are important because they suggest that stress-dependent alignment of the microtubule array is a plausible and sufficient feedback control mechanism to persistently integrate cell geometry and the microtubule-microfibril systems during morphogenesis.

In this paper, we demonstrated how multiple aspects of tensile forces can provide patterning information to orient the microtubule array at cellular scales. We expect this patterning system operates to link any process that changes the stress distributions. This includes tissue-layer interactions that can generate strong tensile forces in the epidermis, as well as geometric and compositional variables that have cell-autonomous effects on cell wall stress (Fig. 5D). These multiscale cellular interactions can affect the magnitude, direction, and anisotropy of stress, and these parameters can operate alone or in combination to influence the microtubule array (Fig. 5D). We predict that this regulatory scheme operates in all plant cells that use a polarized diffuse growth mechanism. This feedback control system would not only enable cells to maintain cell wall integrity by generating oriented fibers in response to high stress, but also enable the cytoplasm to sense cell geometry and program predictable growth outputs.

**Figure 5.**
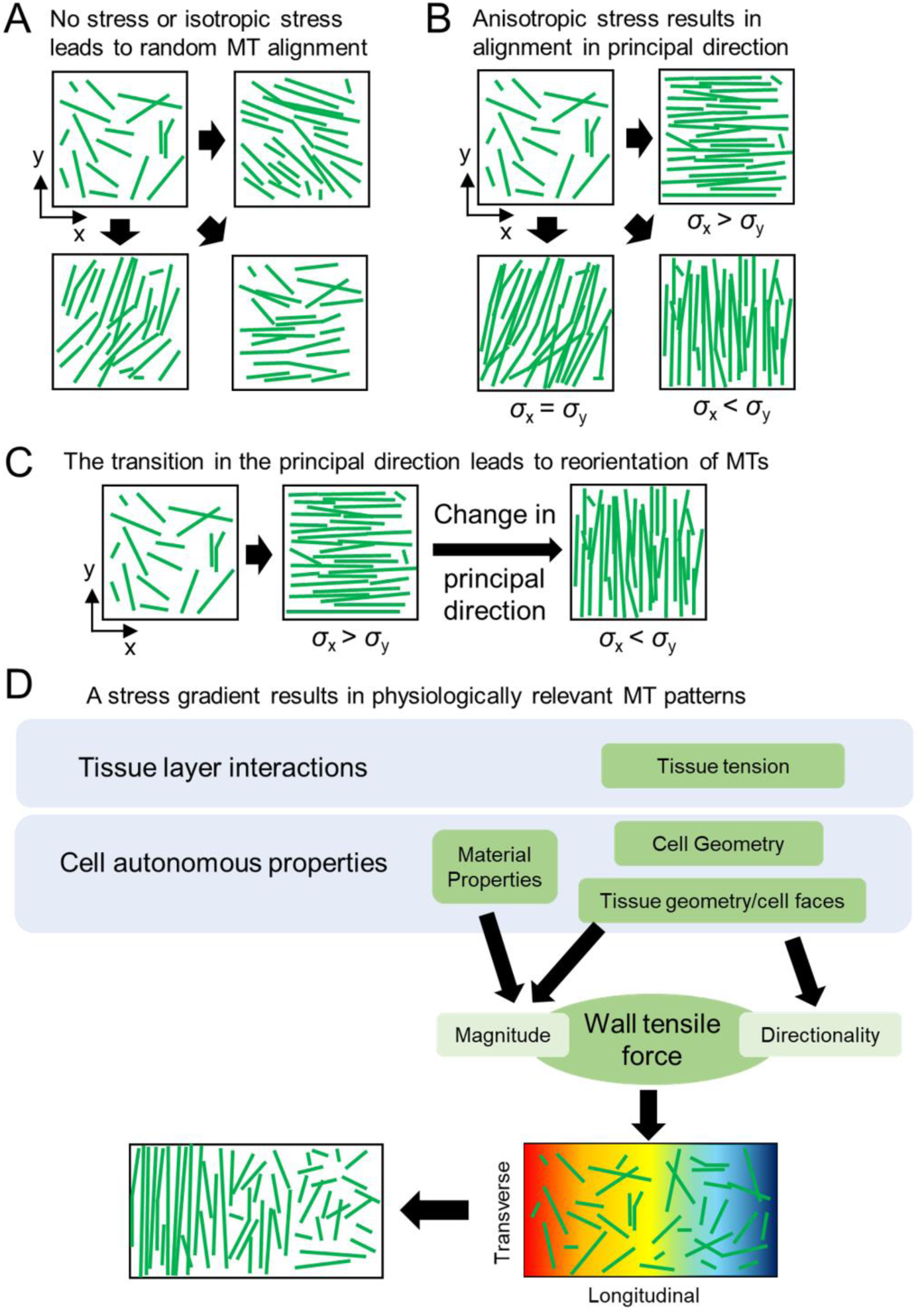
A proposed model for how mechanical intervention biases the microtubule network and leads to its global reorientation. (A) When MTs are not influenced by mechanical stress, the network can form aligned MT bundles in random directions. (B) As the network is subject to anisotropic stress pattern, MTs can align and orient in the direction parallel to the principal stress, to various extents based on which parameter is regulated by stress. (C) Time-dependent transition of the principal direction of network stress can lead to dynamic remodeling and MTs thus globally reorient the MTs. (D) With a biological cell-based stress gradient, MTs can form physiologically relevant patterns with transverse bands near the cell apex during anisotropic diffuse growth, reminiscent of the alignment of MTs parallel to tip-biased, shape-derived, material-property-based anisotropic cell wall stress.

## Acknowledgments

This work was supported by the National Science Foundation MCB grant no. 1715544 to D.B.S and T.Y.K. We greatly appreciate Dr. Joseph Turner and his lab for providing us with the previous modeling data from the finite element analysis of trichome growth. We also want to greatly thank the Turner lab as well as Samuel Belteton from Dr. Szymanski’s lab for insightful discussion about microtubule localization during plant morphogenesis in trichomes and pavement cells.

## Author Contributions

**Conceptualization:** Jing Li, Daniel Szymanski, Taeyoon Kim.

**Methodology:** Jing Li, Taeyoon Kim.

**Investigation:** Jing Li.

**Project Administration:** Daniel Szymanski, Taeyoon Kim.

**Funding Acquisition:** Daniel Szymanski, Taeyoon Kim.

**Formal Analysis:** Jing Li.

**Supervision:** Daniel Szymanski, Taeyoon Kim.

**Writing – Original Draft:** Jing Li, Daniel Szymanski, Taeyoon Kim.

## SUPPLEMENTAL INFORMATION

**Table S1.** Stochastic and deterministic event parameters in addition to dynamic instability.

| Notation | Value | Description |
| --- | --- | --- |
| $\theta_z$ | 40° [1, 2] | Entrainment angle upon collision |
| $f_{nuc}$ | 100 [$\mu\text{m}^{-2} \text{min}^{-1}$] | Nucleation rate |
| $p_{cat}$ | 0.2 - 0.8 | Probability of catastrophe upon collision |
| $p_{zip}$ | 1.0 [2] | Probability of zippering |
| $\delta$ | 0 – 50 nm | Spacing between bundled MTs |
| $p_g$ | 0.65 [3] | Phase of growth % |
| $p_p$ | 0.10 [3] | Phase of pause % |
| $p_s$ | 0.25 [3] | Phase of shorten % |
| $\delta_{seg}$ | 0.1 [$\mu\text{m}$] | MT segment length |

### Stress anisotropy and collision-induced catastrophe coregulate microtubule ordering

We showed earlier that microtubule can form bundles without stress anisotropy (Fig. S2C). Such stress-free alignment is mainly attributed to collision-induced catastrophe. Although zippering, the alignment of two microtubules after collision with small contact angles, can help microtubules align, the reorientation of microtubule bundles can be facilitated only by the collision-induced catastrophe. Microtubules can turn over more frequently in the presence of the collision-induced catastrophe, which helps microtubules align with each other. The collision-induced catastrophe could be more or less important if there is stress anisotropy that can also lead to microtubule ordering. To understand how these two different mechanisms facilitate microtubule alignment in a cooperative or antagonistic manner, we ran simulations with different probabilities for the collision-induced catastrophe and different levels of stress anisotropy. The probability of the catastrophe (*P*_cat_) was varied between 0.2 and 0.8 to be consistent with the previous literature [4]. In cases with a stress-dependent variation in three parameters – the polymerization rate, the depolymerization rate, and the rescue frequency – the low probability of collision-induced catastrophe resulted in nearly randomly oriented microtubules due to substantial increases in cross-over events. As *P*_cat_ increases, the dependence of microtubule ordering on the degree of stress anisotropy became stronger in cases with all the three parameters (polymerization, depolymerization rates and rescue frequency) varied by stress (Fig. S6). The cases with the polymerization rate and the rescue frequency as a stress-sensitive parameter showed almost equal dependence of microtubule ordering on *P*_cat_ and stress anisotropy level (Figs. S6A, D). By contrast, the case with the depolymerization rate varied by stress showed that microtubule ordering is more sensitivity to *P*_cat_ than stress anisotropy (Fig. S6B). At small stress anisotropy level, the order parameter in the case with the depolymerization rate varied by stress significantly increased when *P*_cat_ was changed from 0.6 to 0.8. However, it remained at very small values when *P*_cat_ was below this range, unlike order parameter in the cases of the polymerization rate and the rescue frequency varied by stress showing a consistent increase with an increase in *P*_cat_.

In the case with the catastrophe frequency modulated by stress, microtubules were aligned well even at low *P*_cat_ when intermediate stress anisotropy level was imposed (Fig. S6C). Both stochastic catastrophe and collision-induced catastrophe contribute to the turnover of microtubules. By decreasing the stochastic catastrophe frequency, microtubule ordering became less dependent on stress anisotropy level since the turnover of microtubules are predominantly regulated by collision-induced catastrophe. By contrast, when stochastic catastrophe events take place frequently, microtubule ordering depends more on stress anisotropy because microtubule turnover became more dependent on free catastrophe (Figs. S6E-F).

In sum, the combinative effects of stress anisotropy and collision-induced catastrophe result in microtubule alignment and ordering. Stress anisotropy directly impacts the efficiency of the alignment of microtubules with the direction of principal stress, whereas collision-induced catastrophe controls the portion and turnover of misaligned microtubules.

## Supplemental Figures

**Figure S1.**
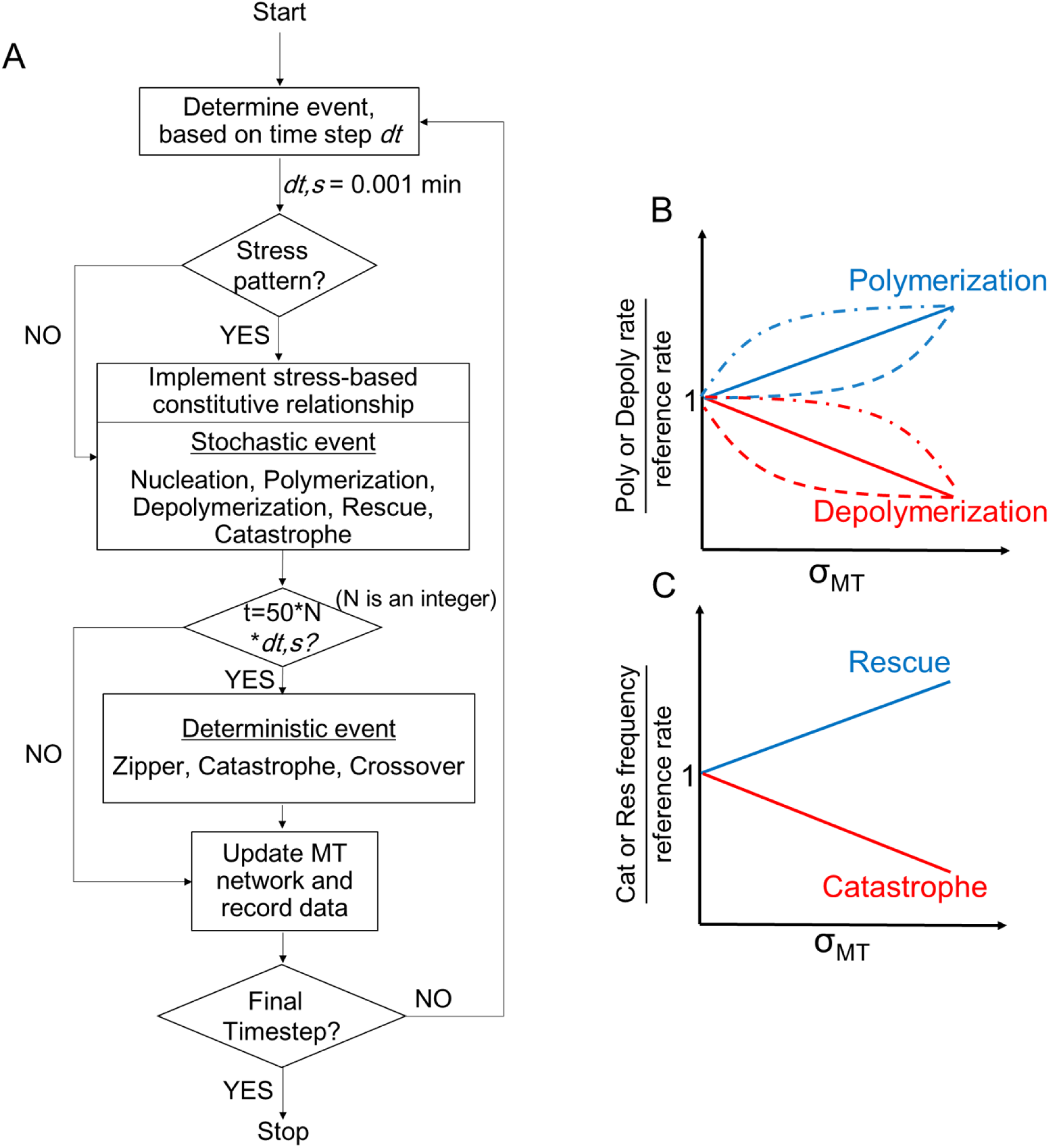
Model description in details. (A) The main flowchart showing the simulation iterative steps. (B-C) The constitutive relationship between plus end dynamics and the stress acting on microtubules. (B) An increase in stress would enhance polymerization rate and suppress depolymerization rate. Solid lines: linear constitutive relationship. Dashed lines: convex or concave constitutive relationship. (C) An increase in stress would enhance rescue frequency and suppress catastrophe frequency (log scale).

**Figure S2.**
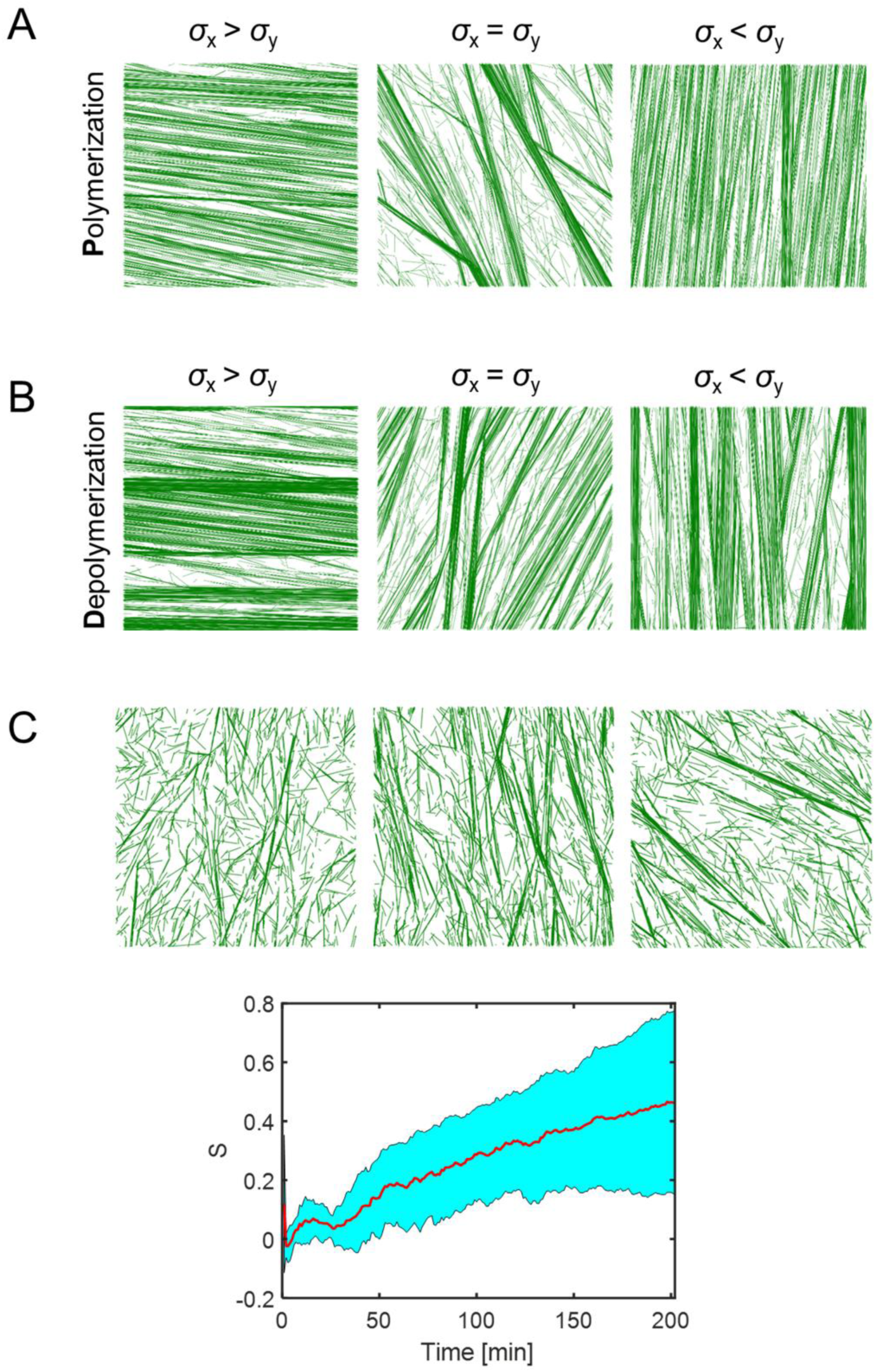
Reference condition and stress-based simulation with larger network domain. (A-B) Steady state MT morphology for network with size of 20 μm x 20 μm. All snapshots are taken at 100 min. MTs are subject to network stress predominant in y direction (right), isotropic (middle), predominant in x direction (left). In A and B, polymerization rate is enhanced and depolymerization rate is suppressed in alignment with principal stress, respectively. (C) Representative cases showing steady state MT morphology for network with size of 10 μm x 10 μm but no stress. All snapshots are taken at 100 min. Time evolution of order parameter (n = 20 cases) with average (red) and standard deviation (cyan area). The order parameter is calculated with respect to a dominant angle.

**Figure S3.**
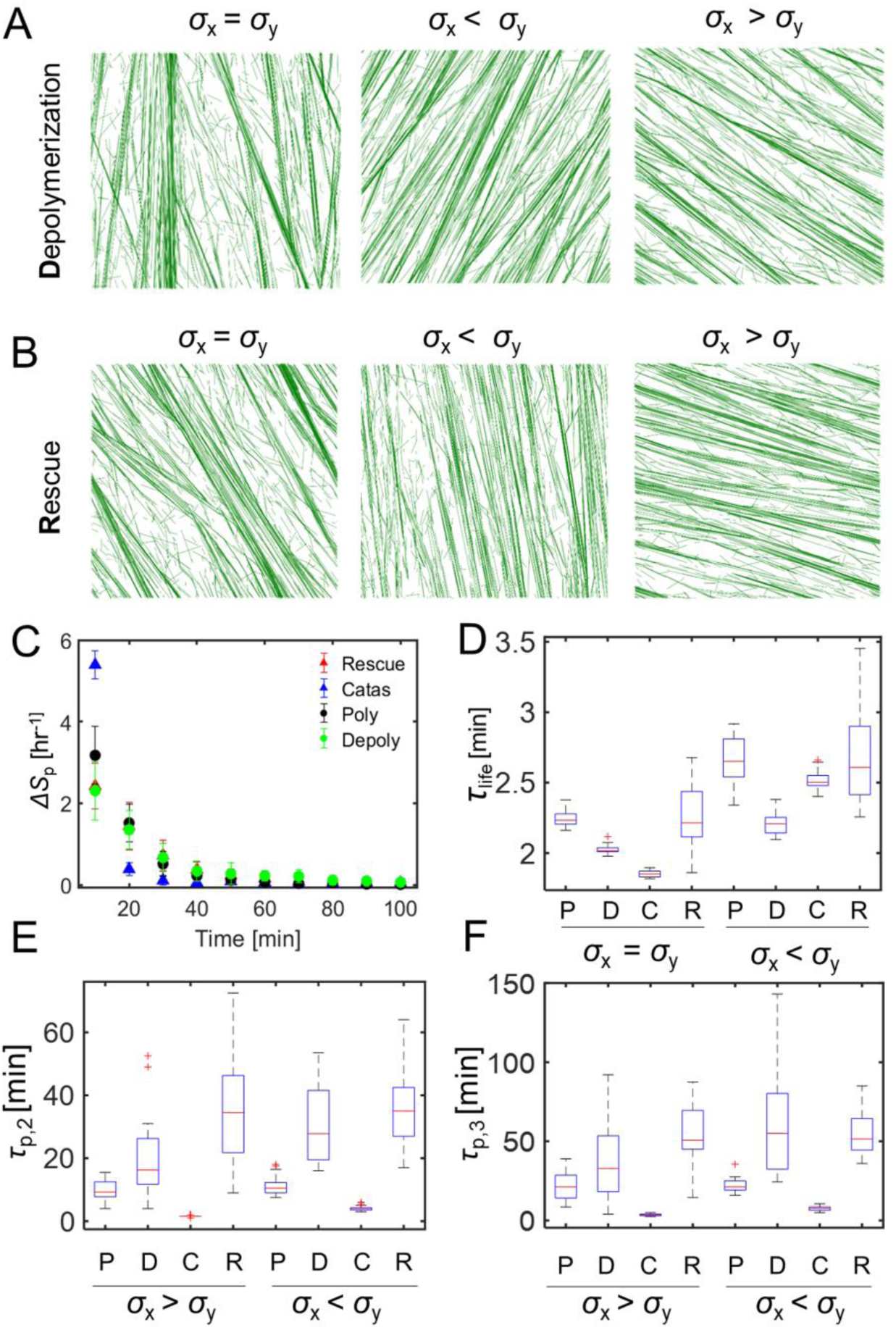
Anisotropic stress regulates network dynamics and microtubule lifetime. (A-B) Steady state MT network morphology. All snapshots are taken at 100 min. MTs are subject to network stress predominant in y direction (right), isotropic (left). In A and B, depolymerization rate is suppressed, and rescue frequency is enhanced in alignment with principal stress, respectively. (C) Time evolution of the rate of change in network order parameter for four different conditions with anisotropic stress. When there is a constitutive relationship between stress and catastrophe frequency, the efficiency of network alignment is significantly increased. In the case of polymerization rate, the increase is smaller. (D) Summary of the average lifetime of microtubule *τ_life_* for all different conditions with isotropic vs. anisotropic stress in which principal stress influences individual stochastic parameter independently. (E-F) Boxplot of the time constants (second and third) acquired from cases with anisotropic stress predominant in x or y directions in all conditions. There is no significant difference due to directional of the principal stress. Data for each condition are averaged over 20 simulations. P: polymerization, D: depolymerization, C: catastrophe, R: rescue.

**Figure S4.**
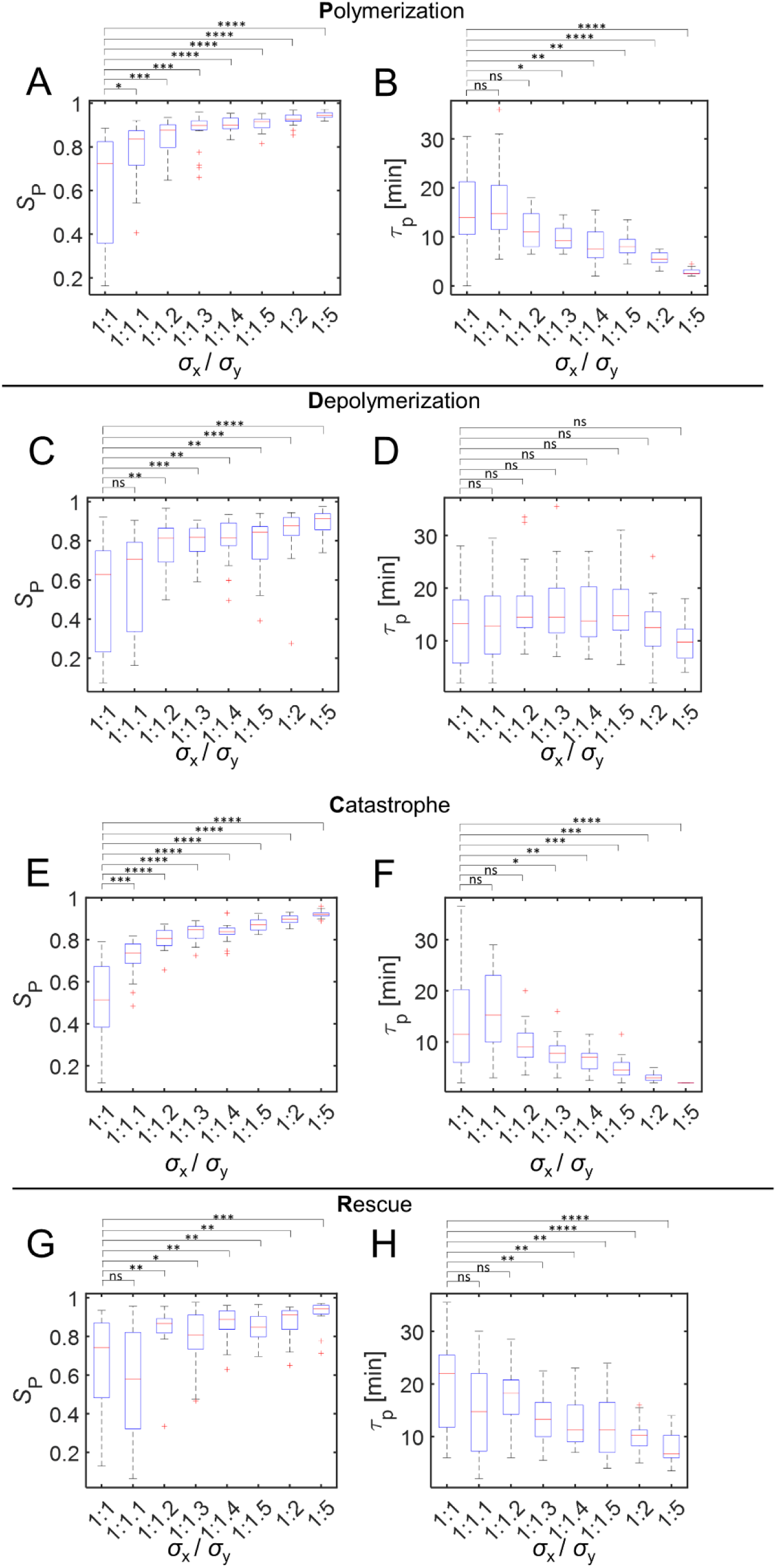
Effect on the microtubule ordering by ratio of anisotropic stress in the network. Isotropic stress has a ratio of 1:1. The anisotropic stress ratio is increased to 1.1, 1.2, 1.3, 1.4, 1.5, 2 and 5. (A, C, E, G) The order parameter of network at steady state as a function of stress anisotropy. (B, D, F, H) The first time constant of network order parameter as a function of stress anisotropy. Boxplot for each order parameter and time constant are acquired from 10 simulations in each case. P: polymerization (A,B), D: depolymerization (C,D), C: catastrophe (E,F), R: rescue (G,H); ns p>0.05, * p<0.05, ** p<0.01, *** p<0.001, **** p<0.0001.

**Figure S5.**
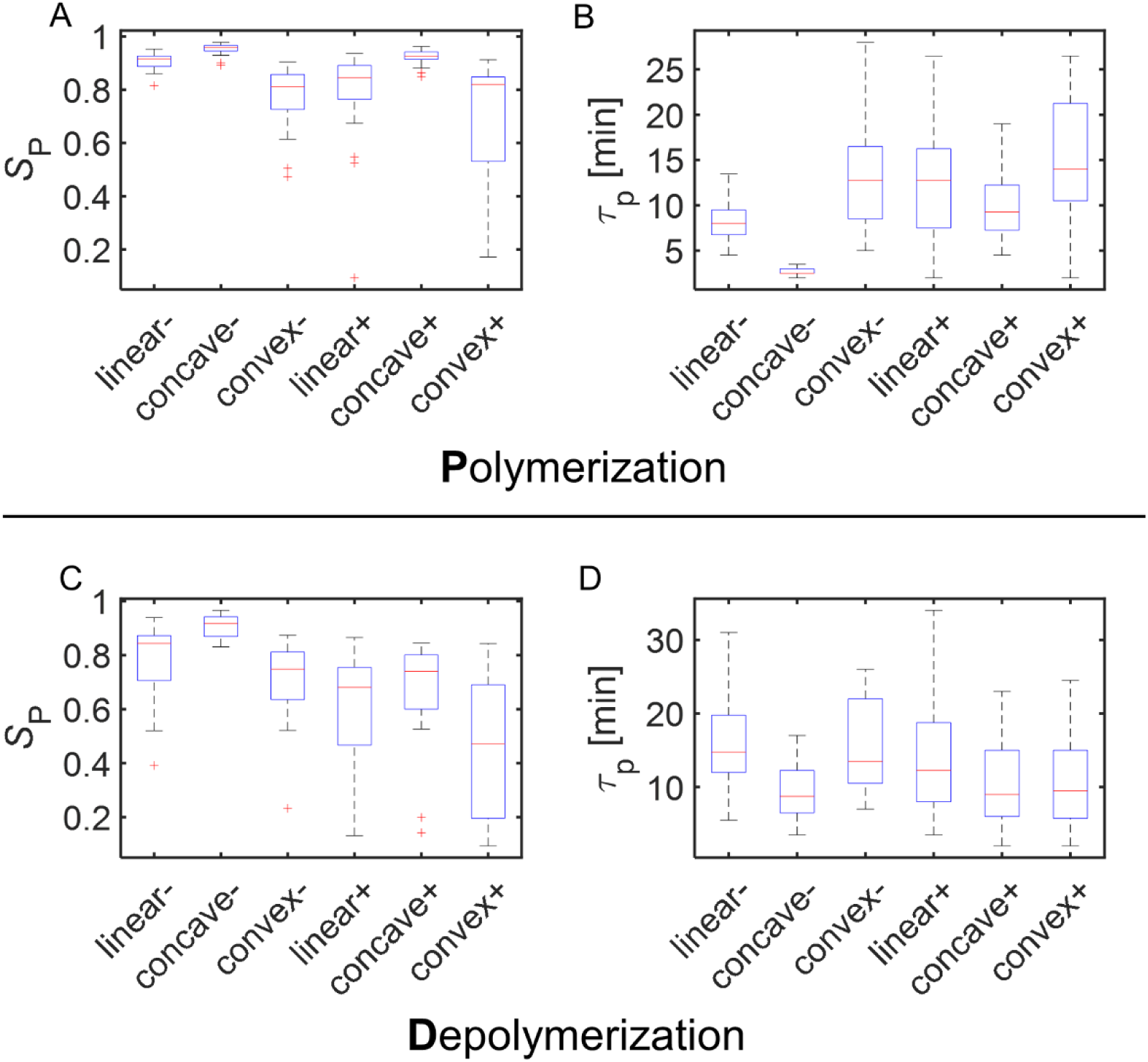
Effect on the microtubule ordering by various types of constitutive equation regulated by stress. The stress anisotropy is maintained at a constant ratio of 2. (A-D) A convex and concave function regulating the stress and polymerization (A, B) rate or depolymerization rate (C, D) is incorporated. Various combinations of functions (linear, concave, convex) and shape (suppress – vs enhance +) are employed. Linear function is a reference condition. Order parameter of network and time constant of increase in order parameter in the case of polymerization (A, B) and depolymerization (C, D) regulated by stress, respectively. A concave function leads to an increase in order parameter at steady state and efficiency of alignment while a convex function inhibits them. n=10 for each condition.

**Figure S6.**
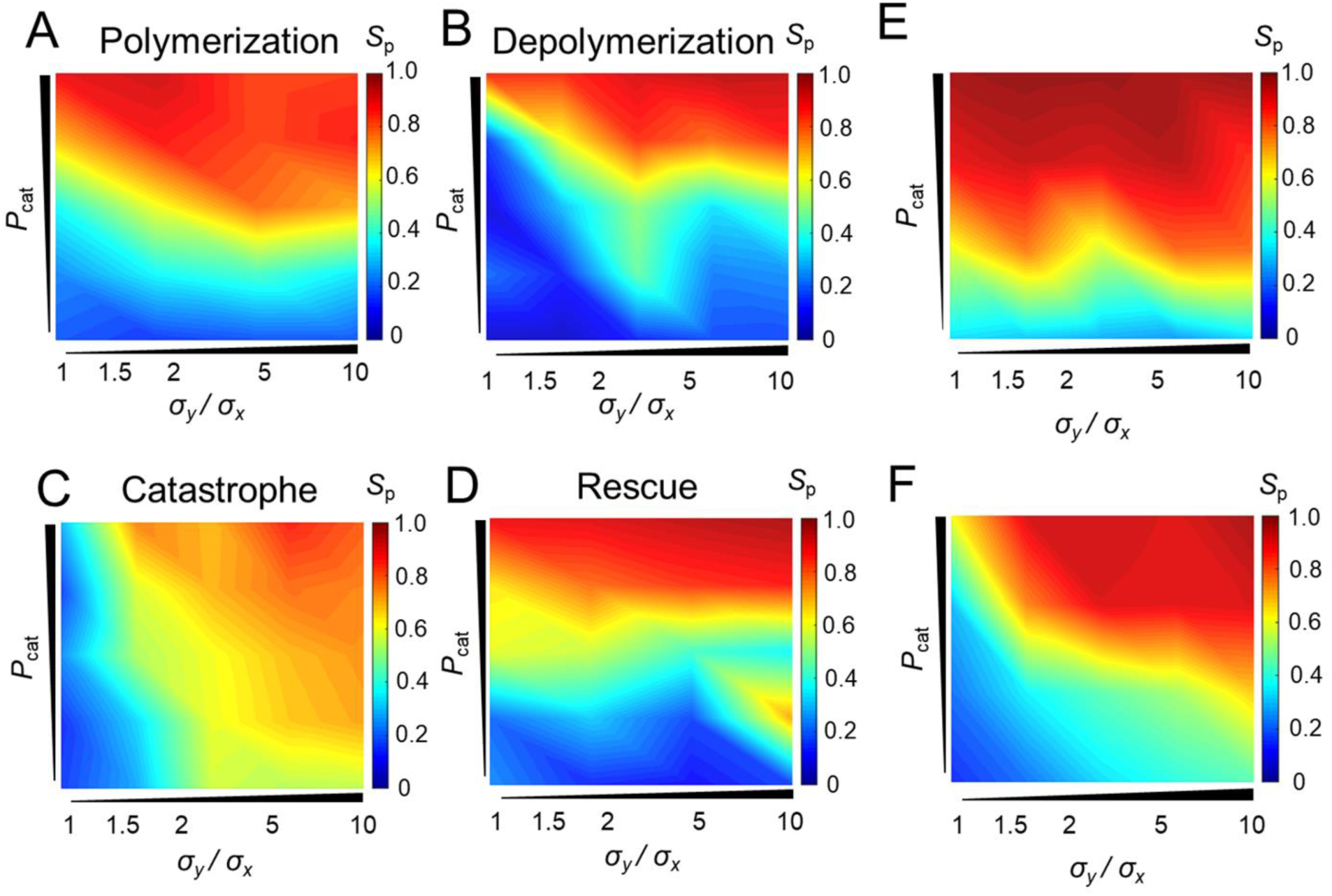
Stress anisotropy and collision-induced catastrophe coregulate microtubule ordering. In the cases when there is a constitutive relationship between stress and stochastic dynamics of microtubule, a two-dimensional parameterization is performed. (A-D) Heatmap summarizing the order parameter of network at steady state for cases of polymerization (A), depolymerization (B), catastrophe frequency (C) and rescue frequency (D) when both stress anisotropy (x-axis) and collision-induced catastrophe probability *P*_cat_ (y-axis) are varied. Stress anisotropy ratio is varied from 1 to 1.2, 1.5, 2, 5, 10 and collision-induced catastrophe is varied from 0.2 to 0.4, 0.5, 0.8. (E-F) Stress anisotropy, collision-induced catastrophe and free catastrophe frequency show a combinative effect on network alignment. (E) Heatmap of network order parameter when base frequency of free catastrophe is elevated. Due to high free catastrophe frequency, the effect by anisotropy of stress (x-axis) is minimized. By increasing the collision-induced catastrophe probability *P*_cat_ (y-axis), the order parameter increases and remains independent of stress anisotropy. (F) Heatmap of network order parameter when base frequency of free catastrophe is reduced. The dependence of order parameter on stress anisotropy and *P*_cat_ are equal. Higher stress anisotropy with higher *P*_cat_ leads to the highest network alignment.

**Figure S7.**
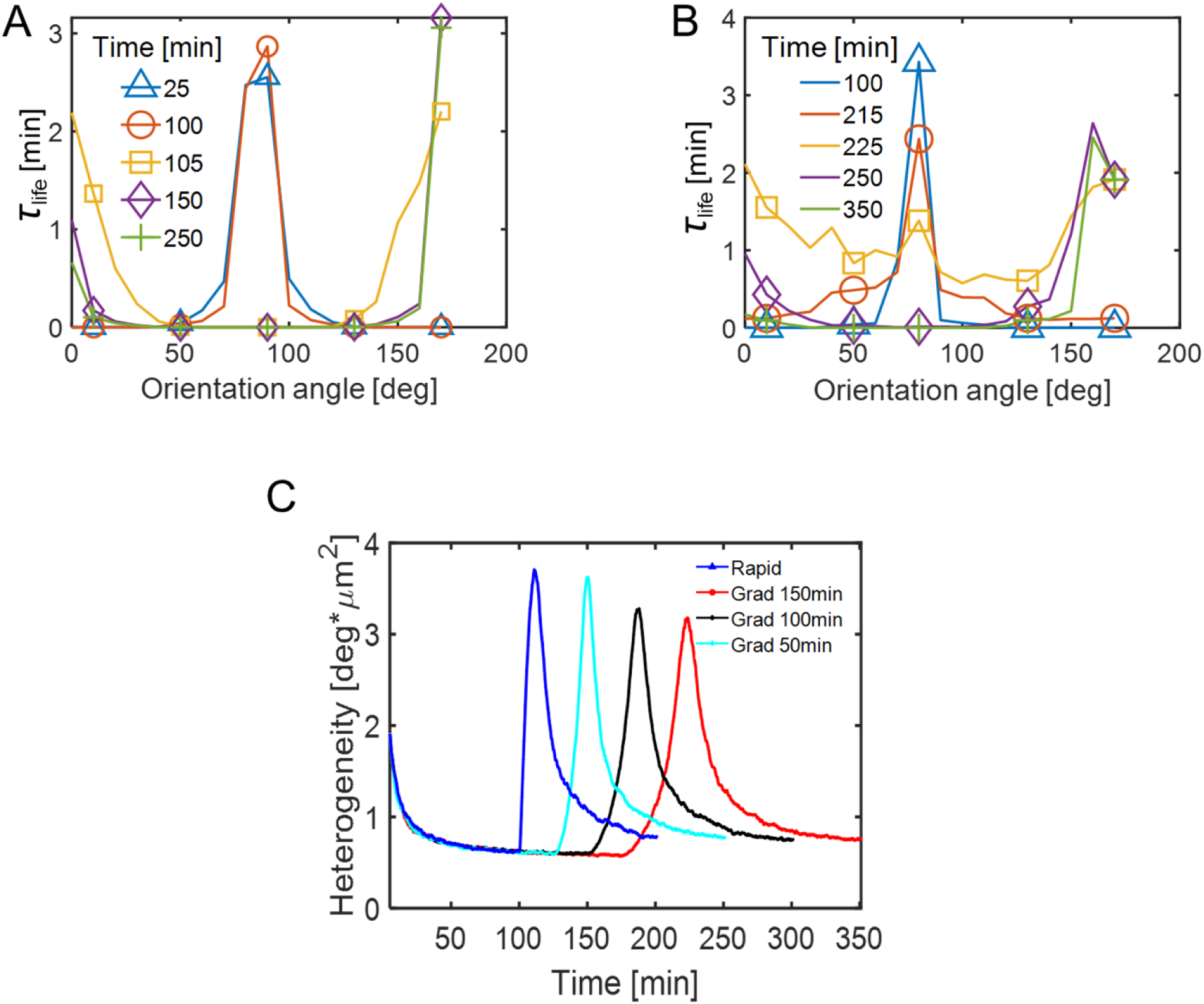
Microtubule dynamics during stress reorientation. (A, B) Distribution of average microtubule lifetime with different orientation angles when stress reorientation occurs rapidly (A) and gradually (B), corresponding to Figures 3A and B. In both cases, the initial orientation of microtubules is predominant near 90° (vertical) with much longer lifetime. After stress pattern reorientation ∼100 min, the microtubules with longer lifetime shift toward 0° or 180° in the horizontal direction until the end of simulation. (C) Network heterogeneity index as a function of time for various stress pattern transition conditions. The heterogeneity is defined as the standard deviation of microtubule angles divided by the total density.

**Figure S8.**
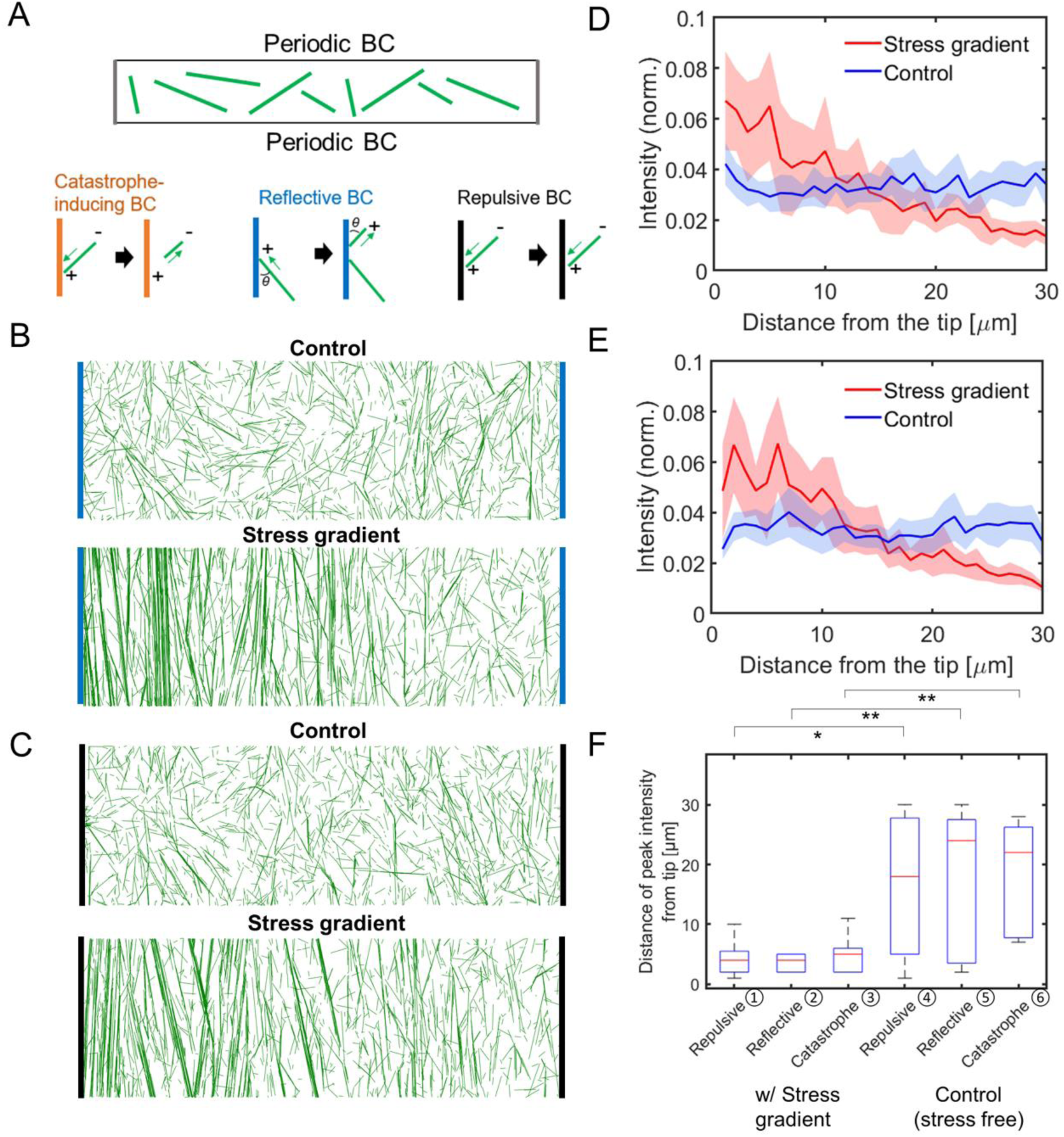
Network with various boundary conditions. (A) Schematic of the network. The long edges are assumed to have periodic boundary conditions. The left and right edges are subjected to three different types of boundary conditions. Catastrophe-inducing boundary is explained in Figure 4. Reflective boundary (blue): when a microtubule encounters a boundary, it keeps growing at the same angle as it hits the boundary in the new direction reflective to the old direction. Repulsive boundary (black): a microtubule hitting a boundary would switch to a pause state. (B, C) Network morphology at 100 min when no stress is included (upper panel) or with a stress gradient (lower panel) for reflective boundaries (B) or repulsive boundaries (C). (D, E) Steady state intensity distribution of microtubules in (B) and (C), with increasing distance from the tip. Without stress, the network is homogeneous. Distinct bundles near the cell tip can form transverse-band-like pattern with a stress gradient regardless of the boundary conditions. (F) Distance of the peak intensity of microtubules from the tip. With a stress gradient, density peaks near the cell tip as transverse bands form. Without stress, the increased homogeneity of network leads to random distribution of local peak intensity.

**Figure S9.**
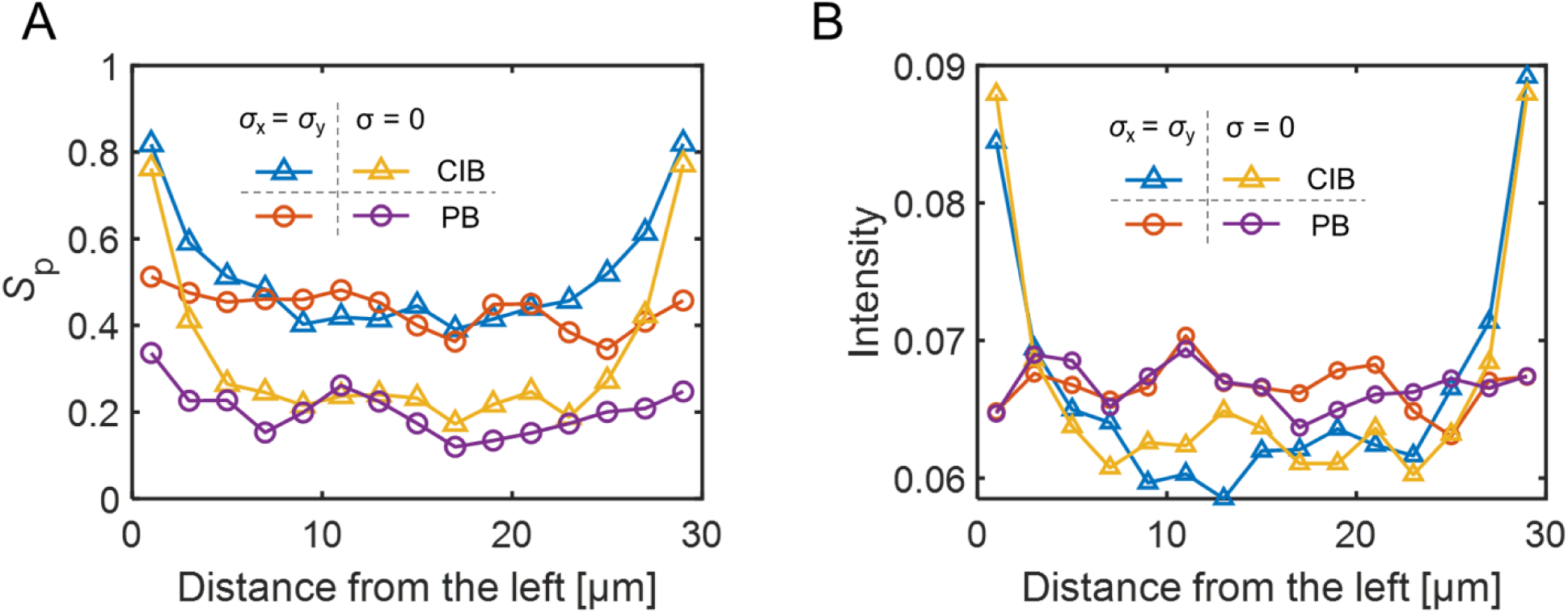
Isotropic stress combined with catastrophe-inducing boundary affects self-organization of microtubules and correlates with local orientation in larger domain. (A) Local order parameter as a function of the distance from the left boundary. (B) Local intensity of microtubules with increasing distance from the left boundary. Legend: Catastrophe-inducing boundary (CIB) with isotropic stress (blue triangle) or no stress (yellow triangle). Periodic boundary (PB) with isotropic stress (orange circle) and no stress (purple circle).

**Figure S10.**
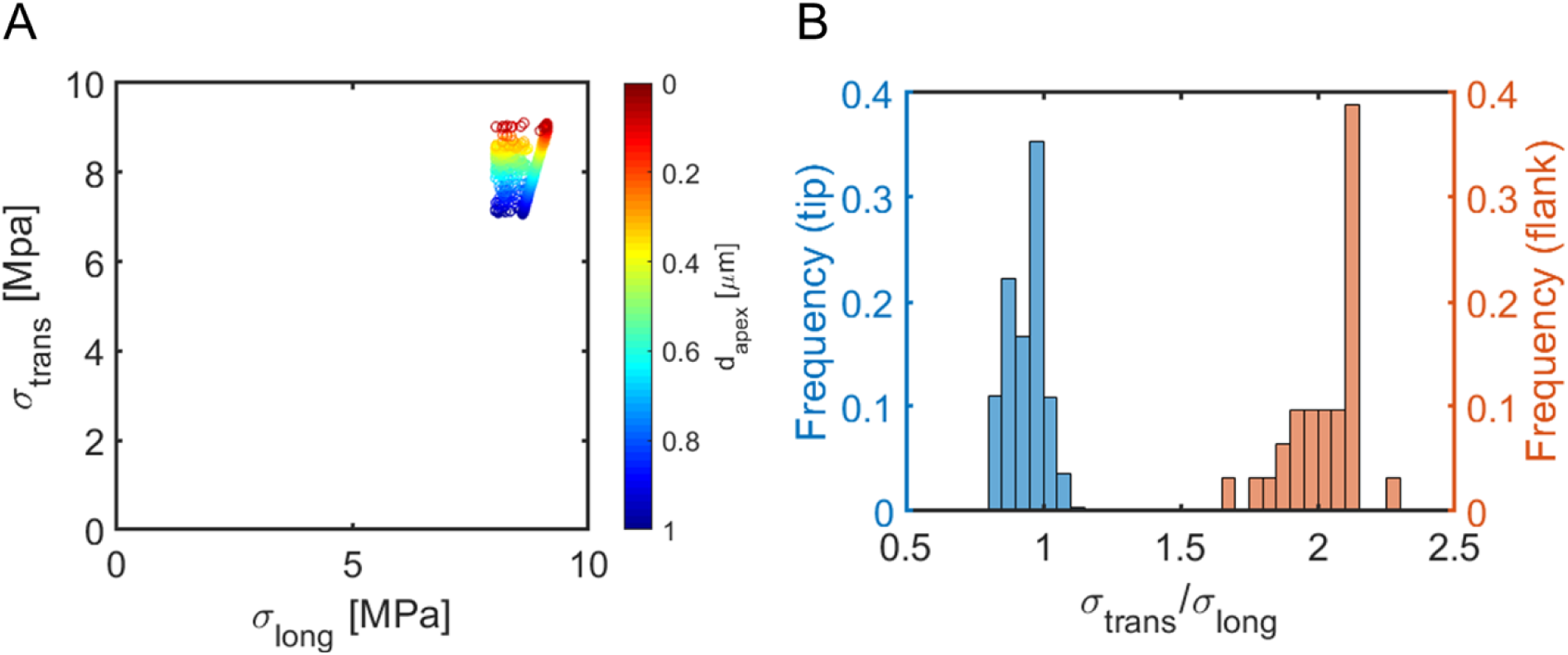
Stress profile from the FEM model. (A) Paired values of longitudinal and transverse stress in the apex of trichome model. Color indicates the distance from apex. (B) Distribution of the stress anisotropy ratio in the tip zone (blue) and flank region (orange). Stress is isotropic in the tip zone while anisotropic in the flank region, as implemented in our simulations.

